# Predatory bacteria impact *C. elegans* life-history traits by modulating microbiota community dynamics and thereby vitamin B12 availability

**DOI:** 10.64898/2026.06.25.734446

**Authors:** Janna Wülbern, Levke Hansen, Catherine Bannon, Manuel Liebeke, Johannes Zimmermann, Brendan J. M. Bohannan, Julia Johnke

**Affiliations:** Evolutionary Ecology and Genetics, Zoological Institute, Kiel University, Kiel, Germany; Max Planck Institute for Marine Microbiology, Bremen, Germany; Department of Metabolomics, Institute for Human Nutrition and Food Science, Kiel University, Germany; Cluster of Excellence Balance of the Microverse, Friedrich Schiller University, Jena, Germany; Institute of Ecology and Evolution, Department of Biology, University of Oregon, Eugene, Oregon, USA

## Abstract

Predatory bacteria such as *Bdellovibrio* are emerging as ecological modulators in microbial communities by restructuring community composition, yet their roles in host-associated microbiomes remain poorly understood. Using *Caenorhabditis elegans* as model host and its defined microbiota, we investigated how two *Bdellovibrio* strains with distinct prey ranges (*B. tiberii* MYbb2 and *B. krueschi* MYbb4) affect microbial community composition and host life-history traits. Both strains consistently altered microbiome composition, with MYbb4 causing more pronounced alpha-diversity shifts and MYbb2 selectively enriching strains of the genus *Ochrobactrum* which coincided with higher host median lifespan. Genome-based predictions indicate that de novo vitamin B12 synthesis by *Ochrobactrum* underlies the observed host phenotype, which was confirmed through quantitative measurements of the vitamin in mono-cell cultures. Employing the *acdh-1*p::GFP transcriptional reporter strain, we confirmed that a diet of B12-producing bacteria suppresses the B12-independent propionate detoxification pathway in the host, demonstrating that bacterially produced B12 is bioavailable to *C. elegans*. Exogenous B12 supplementation assays further confirmed the lifespan-extending effect. Together, these results suggest that predation-driven enrichment of B12-producing bacteria maintains B12 levels sufficient to detoxify propionyl-CoA via the B12-dependent pathway, preventing the accumulation of toxic metabolic byproducts that would otherwise arise under B12-limiting conditions and reduce host lifespan. Our findings demonstrate that predatory bacteria are important drivers of microbiome structure with direct consequences for host physiology, representing an underappreciated ecological mechanism for microbiome modulation.

## Introduction

Virtually all multicellular organisms harbor diverse microbiota, forming a functional unit known as the metaorganism^1,2^, or holobiont^3^. These communities shape host biology by providing nutrients^4,5^, degrading organic compounds^6^, and shaping host development and immunity^7–10^. Microbiota can shape host life-history traits^11^, including reproduction^12^, lifespan^13^, immunity^14^, pathogen resistance^15,16^, host evolutionary trajectories^17,18^, and behavior^8,19^. In *Caenorhabditis elegans*, the microbiota supply nutrients^20^, produce metabolites^21^, and protect against pathogens^22–26^.

Microbiota diversity is expected to be positively correlated with functional redundancy and resilience to disturbance^27^, yet interactions among microbiota members are dynamic and context-dependent, shaped by environmental conditions^28,29^, temporal variation^30^, and by the host itself^31^. Disruptions in microbiota composition have been linked to a range of host diseases^32–34^, from coral bleaching^35^ to inflammatory bowel disease^10,36,37^. Therapeutic interventions, including fecal microbiota transplantations^38^, probiotics^39,40^, or engineered microbial consortia^41^ often result in only transient effects^42,43^. This highlights the need for ecologically informed approaches that treat host-associated microbiota as dynamic ecosystems. One such strategy is introducing a higher trophic level, e.g., predatory bacteria, to restructure and stabilize the microbiota.

Predatory bacteria belonging to the “*Bdellovibrio* and like organisms” (BALOs) are ubiquitous predators that shape microbial community composition across environments, including in hosts^44,45^. These highly motile Gram-negative bacteria prey obligatorily on other Gram-negative bacteria and influence microbiota composition likely through frequency dependent predation (aka kill-the-winner dynamics^46^), whereby numerically dominant prey taxa are selectively reduced, which may promote coexistence with less abundant taxa^47–50^. Most BALOs are periplasmic predators, such as *Bdellovibrio bacteriovorus*, which invade the periplasm of their prey, consume the prey’s cellular contents, and replicate within the host before lysing the prey cell^51^.

BALO presence has been documented in many host-associated microbiota, including zebrafish, stickleback fish^52^, nematodes^53^, arthropods^54^, and mammals (including humans)^55^. There is evidence that BALOs may influence host microbiota diversity and function. For example, a meta-analysis of 16S amplicon sequencing data^56^ positively correlated BALO presence and microbiota diversity across multiple host species, including *C. elegans* and humans, and *Bdellovibrio* DNA has been observed at a higher frequency in healthy human subjects than in patients with microbiota-associated intestinal disease^55^. However, this evidence is all correlative, and experimental tests of the impact of BALOs on host microbiota structure and function are lacking.

To address this, we experimentally manipulated the presence of BALOs in the model animal host, *C. elegans. C. elegans* is a powerful *in vivo* model for studying host-microbe and microbe-microbe interactions. Its native microbiota^53,57,58^ is well-characterized, which led to the development of different standardized laboratory inocula “*C. elegans* microbiota” or “CeMbio” that vary in microbial species richness^59,60^. To investigate how BALO predation influences host-associated microbiota and host function, we inoculated *C. elegans* strain N2 with CeMbio microbiota. We then introduced a single pulse of either a *Bdellovibrio* strain or the same volume of sterile buffer.

We replicated this experiment using two different *Bdellovibrio* species originally isolated from the immediate environment of wild *C. elegans* populations (*B. tiberii* MYbb2 and *B. krueschi* MYbb4)^52^. These species differ in prey range; MYbb2 can feed on multiple CeMbio genera, but MYbb4 can only feed on bacteria of the genus *Ochrobactrum*. We hypothesized that BALO presence should induce changes in microbiota diversity and composition relative to the buffer control, and that these effects should be stronger in response to the BALO species with the broader prey range (MYbb2). Furthermore, we hypothesized that BALO-induced changes in the microbiota would alter microbiota function (such as vitamin production) and host traits (such as lifespan and fertility).

## Material and Methods

### Strains and their maintenance

Resources used in this study and their respective sources are listed along with a short note about their maintenance in Tab. 1. Details regarding media, bacterial cultivation, and worm maintenance are given below.

**Table 1:** Organisms used in this study. Bacterial and worm strains are listed with their respective sources and information on their maintenance conditions.

| Resource | Source | Maintenance |
| --- | --- | --- |
| **Bacterial strains** |  |  |
| **CeMbio(12)** | Dirksen et al., 1016, 2020<sup>57,59</sup> | LB-medium, 42 h, 20°C, 300 rpm |
| **CeMbio(43)** | Johnke et al., 2020; Zimmermann et al., 2024<sup>53,60</sup> | LB-medium, 42 h, 20°C, 300 rpm |
| ***Bdellovibrio tiberii* MYbb2** | Maher et al., 2025<sup>52</sup> | DNB-medium, 2x 24 h, 28°C, 180 rpm |
| ***Bdellovibrio krueschi* MYbb4** | Maher et al., 2025<sup>52</sup> | DNB-medium, 2x 24 h, 28°C, 180 rpm |
| ***Escherichia coli* ML35** | Buttin et al., 1956; Casale et al., 2018<sup>61,62</sup> | LB-medium, 24 h, 28°C, 180 rpm |
| ***Ochrobactrum* sp. MYb248** | Dirksen et al., 2016<sup>57</sup> | LB-medium, 24 h, 28°C, 180 rpm |
| **Worm strains** |  |  |
| ***Caenorhabditis elegans* strain N2** | Brenner, 1974<sup>63</sup> and provided by the CGC | V-medium, 20°C, 300 rpm |
| ***C. elegans* strain VL749**<br>**Genotype: *wwls24[acdH-1p::gfp + unc-119(+)]*** | Arda et al., 2010; MacNeil et al., 2013<sup>64,65</sup> and provided by the CGC | NGM agar, 20°C |

#### *C. elegans* maintenance

The transgenic *C. elegans* strain *acdh-1*p::gfp was maintained on nematode growth medium (NGM) agar plates seeded with the standard laboratory *E. coli* strain OP50 at 20°C according to standard procedure^66^.

Similarly, worm populations of wildtype *C. elegans* strain N2 were grown according to the standard procedure^66^ and frozen at -80°C. Freshly thawed worms were revived and passaged for two generations in viscous medium (see below) seeded with CeMbio microbiota at 20°C for two generations (6 days).

Before each experiment, age synchronization was achieved by bleaching gravid hermaphrodites with an alkaline hypochlorite solution. Eggs were incubated in S-buffer (100 mM NaCl, 6.5 mM K_2_HPO_4_, 43.5 mM KH_2_PO_4_) overnight on a shaker.

#### CeMbio microbiota maintenance

The representative community of natural *C. elegans* microbiota, CeMbio(12)^59^, comprising 12 bacterial strains (Tab. 2) served as food sources for *C. elegans* N2.

**Table 2:** CeMbio strain identities and prey range of MYbb2 and MYbb4.

| ID | Species | Predation |  |
| --- | --- | --- | --- |
|  |  | *B. tiberii* MYbb2 | *B. krueschi* MYbb4 |
| **MYb10** | *Acinetobacter guillouiae* | **TRUE** | FALSE |
| **JUb44** | *Chryseobacterium scophthalmum* | FALSE | FALSE |
| **BIGb0172** | *Comamonas piscis* | FALSE | FALSE |
| **BIGb0393** | *Pantoea nemavictus* | **TRUE** | FALSE |
| **CEent1** | *Enterobacter cloacae* | **TRUE** | FALSE |
| **JUb66** | *Lelliottia amnigena* | **TRUE** | FALSE |
| **MYb71** | *Ochrobactrum vermis* | FALSE | **TRUE** |
| **MYb11** | *Pseudomonas lurida* | FALSE | FALSE |
| **MSPm1** | *Pseudomonas berkeleyensis* | FALSE | FALSE |
| **BIGb0170** | *Sphingobacterium multivorum* | FALSE | FALSE |
| **JUb134** | *Sphingomonas molluscorum* | FALSE | FALSE |
| **JUb19** | *Stenotrophomonas indicatrix* | FALSE | FALSE |

CeMbio microbiota cultivation and inoculum preparation were conducted as previously described^60^ with minor adjustments. In short, all 12 bacterial strains were individually incubated in 4 ml lysogeny broth (LB) using 48-well 6 ml polypropylene storage plates for 42 h at 20°C on a circular vibrating shaker at 350 rpm. To ensure similar cell numbers per strain, 0.2-2 ml of the 12 individual bacterial strains were combined and washed thrice in sterile S-buffer for inoculum preparation.

#### BALO strains MYbb2 and MYbb4 maintenance

Predatory bacteria *B. tiberii* MYbb2 and *B. krueschi* MYbb4^52^ were used in all BALO-related experiments.

MYbb2 and MYbb4 were co-cultivated for maintenance with either *E. coli* ML35 or *Ochrobactrum* sp. MYb248, respectively. Prey cells ML35 and MYb248 overnight cultures were grown in LB medium, continuously shaking for 16 hours at 28°C and 180 rpm. BALO strains were separately maintained for up to one month on double-layered diluted nutrient broth (DNB) agar plates (1:10 diluted nutrient broth, supplemented with 2 mM CaCl_2_ and 3 mM MgCl_2_). The top layer contains respective prey cells and BALO plaque-forming units (PFUs) according to standard protocol^67^.

For BALO cultivation in liquid medium, we followed the protocol of Remy *et al.*, 2022^68^ with minor adaptations. In short, single plaques were aspirated from double-layered DNB (1:10 diluted nutrient broth, 2 mM CaCl_2_, 3 mM MgCl_2_) agar plates, released into a sterile glass tube with 2.7 ml DNB medium with 300 µl of *E. coli* ML35 (final OD_600_ =1), and incubated overnight shaking (180 rpm) at 28°C. After 24 hours, 100 µl of the co-culture was passaged to fresh 2.6 ml DNB medium with 300 µl of *E. coli* ML35 (final OD_600_ =1) and incubated overnight shaking (180 rpm) at 28°C. Before each experiment, BALO cells were separated from prey cells by filtration through 0.45 µm filters. For quantification, we followed the protocol of a SYBR Green (Thermo Fisher Scientific, #S7563) quantification assay^68^. Simultaneously, double-layered DNB agar plates were prepared according to standard protocol^67^ to verify SYBR Green assay results. Ultimately, a single pulse of 10^5^ –10^7^ BALO cells/ml was added at the start of each experiment.

### Determination of BALO prey range

We tested all 12 strains from the CeMbio to determine the prey range of both BALO strains (Tab. 2). Prey cell overnight cultures were grown in LB medium, continuously shaking for 16 hours at 28°C and 180 rpm. Predation was tested in both (i) liquid medium and on (ii) plaque assay plates. For BALO cultivation in (i) liquid medium, we followed the protocol of Remy et al.^68^ with minor adaptations, as described above. For each prey strain, three biological replicates (single plaques) of BALOs were tested in two independent runs. Here, prey clearance was used as a proxy for predation success, hence, co-cultures were monitored for turbidity decrease and checked for cell activity microscopically. After 48 h of incubation, BALO cells were separated from prey cells by filtration through 0.45 µm filters. For plaque-assays, we cultivated both BALO strains with their respective maintenance prey (as described above), to ensure sufficient cell density. Then, (ii) double-layered DNB agar plates were prepared according to standard protocol^67^ for each replicate with respective prey strain. Plates were incubated for 5-7 days. Appearance of plaques indicated successful predation.

### Longitudinal experiments

Experiments were conducted in viscous medium (V-medium), based on Papkou et al. (2019)^69^. V-medium is a modified S-buffer supplemented with a non-toxic thickening agent, hydroxypropyl methyl cellulose (HPMC, 10 g/l), salts (3 mM MgCl_2_, 2 mM CaCl_2_, 3 mM MgSO_4_, 1 M KCO_3_), cholesterol (5 µg/ml), and nutrient broth (0.5 g/l). This allowed optimal conditions for *C. elegans* fitness, bacterial dispersion, and BALO predation.

To relate BALO presence to changes in microbiota community dynamics, worm populations co-cultivated with bacterial inocula consisting of the 12 CeMbio strains were exposed to BALO strains MYbb2, MYbb4, or buffer (control) for up to 21 days in independent runs.

Before each run, worms were pre-adapted to the given experimental conditions by cultivation in cell culture flasks (Sarstedt, #83.3911.502), containing V-medium seeded with CeMbio for two generations (6 days). As described above, worm populations were then synchronized by bleaching with an alkaline hypochlorite solution. About 1000 sterile L1 larvae were added to each treatment flask, containing 20 ml V-medium inoculated with CeMbio (final OD_600_ = 1). BALO treatments received a single pulse of MYbb2 or MYbb4 cells (10^5^–10^7^ PFU/ml) at the beginning of the experiment.

Temporal shifts in microbiota composition were assessed across six replicates and three to four time points per treatment for both the worm populations (“worm”) and the directly associated V-medium environment (“medium”). Before sampling, at each time point (day 3, 6, 9, 12, 15, 18, 21), 2 ml of V-medium containing worms of different stages and microbiota were transferred to fresh V-medium to prevent worm overgrowth and nutrient depletion. To assess the microbiota composition of both worm and medium, another 2 ml V-medium was removed from culture flasks, transferred to Falcon tubes (15 ml) diluted with 5 ml PBS, and centrifuged for 1 min at 3500 rpm. At each sampling point, all six biological replicates per treatment were collected.

#### Worm sample processing

Worms were collected from the tube bottom by pipetting and transferred to M9 buffer (3g/l KH_2_PO_4_, 6g/l Na_2_HPO_4_, 5g/l NaCl) supplemented with 0.05% Triton X-100 (M9-T buffer) in Eppendorf tubes. Worm samples were washed following a modified surface bleaching protocol^59^. In detail, worms were washed five times in M9-T buffer by gravity separation. Worms were immobilized prior to hypochlorite washing, by adding 100 µl 10 mM tetramisole to minimize the loss of gut-associated microbiota during washing. By incubating worms in 2% alkaline hypochlorite solution for 2 min, cuticle-associated microbiota was removed. Washing was repeated three times in PBS-T (PBS with 0.05% Triton X-100). Then, 20 µl of PBS-T containing five washed, adult worms were collected in single wells of 96-well plates containing 10 µl Tris-EDTA buffer (pH 8) with 3 mg/µl proteinase K and 3-5 zirconium beads (1 mm) and stored at – 80 °C for at least 16 h.

#### Medium sample processing

After removing the worms from the Falcon tube, the remaining medium, representing the environment of sampled worms, was centrifuged for 15 min at 3500 rpm. The supernatant was discarded, and the pellet was resuspended in 1 ml PBS and collected in a 1.5 ml Eppendorf tube before centrifugation for 5 min at 8000 rpm. The pellets were frozen at -20°C.

### DNA extraction and 16S rRNA amplicon sequencing

#### Worm DNA extraction

Bacterial DNA from worms was extracted, as described previously^59,60^. In short, 96-well plates with frozen worm samples were briefly thawed before disruption for 3 min at 30 Hz in a sample homogenizer (Bead Ruptor 96, Omni International) and subsequent proteinase K digestion (1 h at 55°C, 20 min at 98°C).

#### DNA extraction of medium samples

After thawing microbiota samples from the worm-associated environment (medium), pellets were resuspended in 60 µl PBS-T. From this, 30 µl of bacterial solution was transferred to 96-well plates. A mock community dilution series (10^0^-10^-5^) (ZymoBIOMICS Microbial Community Standard, LOT: ZRC190633) and negative controls (2x TE buffer) were included on each plate. DNA was extracted using the Macherey-Nagel NucleoSpin 96 Tissue kit with 96-well plates and a vacuum manifold with minor modifications: Lysis of microbiota samples was achieved by incubation for 24 h. Extracted DNA was eluted in 80 µl elution buffer (kept at 70°C). Microbiota DNA was stored at -20°C until further processing.

#### Bdellovibrio PCR and qPCR

To confirm *Bdellovibrio* presence prior to 16S rRNA sequencing, bacterial pellets of randomly picked medium samples across different time points and replicates were selected. They were either submitted to diagnostic polymerase chain reaction (PCR) or qPCR, as described below.

Presence of *Bdellovibrio* DNA was verified in medium samples via PCR using primers Bd529F (5ʹ-GTATCGGGAACGTATTCACCG-3ʹ) and Bd1007R (5ʹ-CGGTTGCGCTCGTTGCGG-3ʹ)^70^. Reactions were run under the following cycling conditions: 95°C for 2 min; 36 cycles of 95°C for 30 s, 51°C for 30 s, and 72°C for 30 s; followed by a final extension at 72°C for 5 min. Amplification products were visualized via agarose gel electrophoresis.

Alternatively, the presence of *Bdellovibrio* DNA was checked using a diagnostic quantitative PCR (qPCR)^71^. Amplification of the *Bdellovibrionaceae*16S rRNA gene was performed via one-step PCR using primers Bd347F (5ʹ-GGAGGCAGCAGTAGGGAATA-3ʹ) and Bd549R (5ʹ-GCTAGGATCCCTCGTCTTACC-3ʹ)^70,71^ and the Platinum SYBR Green qPCR SuperMix-UDG (Invitrogen, #11733046). Samples were tested in triplicates. Amplification was performed following the manufacturer’s instructions (UDG incubation for 2 min at 50°C, initial step for 2 min at 95°C and 45 cycles of 15 s at 95°C, 1 min at 60°C, and 30 s at 72°C).

#### 16S rRNA amplicon sequencing

Amplification of the V3V4 hypervariable regions of the bacterial 16S rRNA gene was performed via PCR using barcoded primers 341F (5ʹ-CCTACGGGAGGCAGCAG-3ʹ) and 806R (5ʹ-GGACTACHVGGGTWTCTAAT-3ʹ)^72,73^. A final volume of 26 µl comprised of 5 µl 5X Phusion HF buffer, 0.5 µl dNTP (10 mM), 0.3 µl Phusion Hot Start II Polymerase (2 U/µl; Thermo Fisher Scientific), 9.2 H_2_O, 4 µl of each primer (100 µM), and 3 µl DNA template were prepared. A no-template control (NTC) and a mock community DNA standard (ZymoBIOMICS Microbial Community Standard, LOT: ZRC190633) were included. The resulting PCR products were purified and normalized using the SequalPrep Normalization Plate kit (Thermo Fisher Scientific) according to the manufacturer’s instructions. Final equimolar libraries were sequenced using the paired-end MiSeq reagent kit v3 (2 × 300 bp chemistry) on the MiSeq platform (Illumina Inc.).

### Microbiome 16S rRNA data analysis and statistics

Bioinformatic analysis was performed using the DADA2 pipeline. Sequence quality control and denoising were initially conducted separately for each sequencing run to generate run-specific error models. Following denoising, the resulting sequence tables were merged into a single global table. Taxonomic assignment was then performed on this combined dataset, ensuring that Amplicon Sequence Variants (ASVs) were consistently identified and named across all experiments. In detail, primers were trimmed from the raw MiSeq sequences using cutadapt (v2.6)^74^. Sequences were processed using DADA2 (v1.34.0)^75^ and R (v4.4.1)^76^. Reads were filtered and trimmed using standard filtering parameters (maxN=0, truncQ=2, rm.phix=TRUE and maxEE=c(2,2)) with truncation lengths set to 260 (forward) and 180 (MYbb2 experiment reads) or 190 bp (MYbb4 experiment reads) for reverse reads. Sequence tables were then merged and taxonomy was assigned using the assignTaxonomy and addSpecies commands and the SILVA database (v138.2). These sequences were then aggregated^60^ by aligning them to a blast database containing the reference 16S sequences from the respective CeMbio community genomes generated with makeblastdb^77^. Alignment was done using BLASTn. To ensure high-confidence taxonomic assignments, we set parameters to >99.5% sequence identity and >95% query coverage.

As copy number variation of the 16S rRNA gene may bias abundance estimates^78^, we applied copy number normalization prior to analyzing the CeMbio data by dividing each ASV’s abundance by its corresponding copy number. For these strains closed genomes are available^59^, providing information on 16S rRNA gene copy number variation.

Microbiome sequence data were then analyzed with phyloseq^79^ (v1.48.0). Sequences assigned to chloroplasts (Class: Chloroplast) or mitochondria (Family: Mitochondria) were removed. For downstream analyses, samples with fewer than 1,000 total reads were excluded.

#### Diversity analysis

ASV counts were rounded to integers prior to diversity estimation. Alpha diversity was estimated using observed richness, as implemented in the estimate_richness function of phyloseq by vegan^80^. To model changes in observed richness over time, a generalized linear mixed model (GLMM) with a generalized Poisson distribution was fitted using the glmmTMB package^81^, with treatment, sampling day, and their interaction as fixed effects, and biological replicate as a random intercept. Residual diagnostics were performed using the DHARMa package^82^. Estimated marginal means and pairwise contrasts between treatments were computed using the emmeans package^83^.

Beta diversity was assessed using Aitchison distance, computed as the Euclidean distance on centered log-ratio (CLR)-transformed abundances using the microbiome package^84^. Ordination was performed using redundancy analysis (RDA) as implemented in phyloseq and visualized per sampling day. Differences in community composition between treatments were tested using PERMANOVA (adonis2, vegan, 999 permutations), with biological replicate used as a stratification variable to account for the repeated-measures structure of the data. PERMANOVA was performed both across all time points jointly and separately for each sampling day.

#### Community-weighted vitamin production

Analyses were restricted to worm samples from biological replicates 2, 4, and 6, which corresponded to the flasks used for life-history analyses. Samples were normalized to relative abundances.

To estimate the community-weighted vitamin production potential, relative abundances were aggregated at the CeMbio strain level per biological replicate and treatment at the final sampling time point of each experiment (day 21 for MYbb2; day 15 for MYbb4). Experimentally measured mean vitamin concentrations (B1, B2, B12-OH, B6-xine, and others) were obtained from in vitro quantification of individual CeMbio strains (see below). For each strain, the relative abundance was multiplied by the corresponding mean vitamin concentration to yield a weighted contribution. Weighted contributions were summed across all strains per biological replicate and treatment to obtain a community-level weighted vitamin production estimate. Differences between treatments were tested using Wilcoxon rank-sum tests with Benjamini-Hochberg correction for multiple comparisons, implemented in the rstatix package^85^.

#### B12-to-propionate ratio

To assess the balance between B12 production and propionate fermentation within the community, strains were classified either based on the results of the in vitro vitamin measurements (B12 producer: TRUE/FALSE) or genome-inferred pathway information (propionate fermenter: TRUE/FALSE; see below). For each biological replicate and treatment, the summed relative abundance of B12-producing strains was divided by the summed relative abundance of propionate-fermenting strains, yielding a community-level B12-to-propionate ratio. This ratio was calculated at the final sampling time point of each experiment.

#### Genome-based inference of metabolic pathways

Metabolic pathways from all species were predicted using gapseq v1.1 (3fa038c; Sequence DB 1139b8e)^86^. For sequence alignment, we used a bitscore cutoff of 150 and considered pathways to be present if 80% of the enzymes were present or, alternatively, 66% if all key enzymes were found, as done in our previous analysis^87^.

#### Differential abundance analysis

Differential abundance of CeMbio strains between treatments was assessed using ANCOMBC2^88^ to account for compositional bias. Prior to ANCOMBC2 analysis, technical replicates were collapsed to biological replicates (replicates 2, 4, and 6 on the last sampling day). ANCOMBC2 was run with treatment as the fixed effect, Benjamini-Hochberg p-value adjustment, and a prevalence cutoff of 10%.

Rank abundance distributions were computed from mean relative abundances per treatment.

#### Effect size and power analysis

For MYb71 specifically, effect size was quantified using Cohen’s d and Hedges’ g (effsize package^89^), computed from relative abundances across biological replicates (replicates 2, 4, and 6 on the last sampling day). Statistical power was estimated post-hoc using a two-sample t-test framework (pwr package^90^) at α = 0.05, with the observed Cohen’s d as the effect size estimate.

Data visualization was performed using the ggplot2 package (v3.5.1)^91^ unless stated differently. Throughout the statistical analyses in this study, p-values were annotated using asterisks to indicate levels of significance: *** for *p <* 0.001, ** for *p <* 0.01, * for *p <* 0.05, and “ns” for non-significant results (*p >* 0.5).

### *C. elegans* N2 life-history trait assay

Following the longitudinal study on the effect of BALO cells on CeMbio community dynamics, worm populations were evaluated for life-history traits, *i.e.,* lifespan and fertility. After the final sampling point, single L4 larvae were isolated from the remaining material of three randomly selected biological replicates per treatment and briefly surface-washed on empty 3 cm plates with M9-T. Within each replicate, 22 technical replicates were cultured in parallel treatment wells to account for within-replicate variation, unless stated differently. Individual worms were then transferred to single wells of 24-well plates (Sarstedt, #83.3922), each containing 1 ml V-medium inoculated with the respective CeMbio community (final OD_600_ =0.1). To avoid nutrient depletion, alive adults were transferred daily to fresh plates seeded with the respective CeMbio community (final OD_600_ =0.1) until the end of the egg-laying period, with offspring counts and survival scored daily until day of death. Survival was visualized using Kaplan-Meier curves, all statistical analyses were conducted with a Cox proportional hazards mixed-effects model, using the coxme package^92^. In this model, treatment was included as fixed effect and biological replicate as a random intercept to account for repeated measurements across independent plates.

To assess the BALO effect on worm fertility, mean total offspring counts per worm were calculated for each biological replicate (n=3 per treatment) across technical replicates using the wilcox.test() function from the base stats package. Treatment differences were assessed using a two-tailed Wilcoxon rank-sum test (Mann–Whitney U test).

### Validation of bacterial vitamin B12 production

Bacterial vitamin B12 production was tested by expanding upon previous work^25^ and for the 12 members of the CeMbio community using the dietary sensor *C. elegans* strain *acdh-1*p::gfp^65^. In the presence of B12, the expression of the acyl-CoA dehydrogenase-encoding gene *acdh1* (propionate shunt) is downregulated, resulting in the absence of a GFP signal^21,65,93^. Synchronized L1 larvae were grown on PFM (peptone-free medium) agar plates seeded with lawns of *E. coli* OP50 or single CeMbio isolates. Young adults (24 h post L4) were then paralyzed with 10 mM tetramisole and placed onto 2% agarose patches on microscopic slides. Imaging was performed with a Leica stereomicroscope (M205 FA, Wetzlar, Germany). For comparability of representative images, the same magnification and exposure time for the fluorophore (GFP) signal settings were used (BF = 20, GFP3 = 300, intensity (K) = 2400, zoom = 8x) while contrast and brightness were adjusted.

### B-vitamin quantification by LC-MS

The biosynthetic potential of the CeMbio microbiota to produce B-vitamins was evaluated using targeted quantification of hydroxo-cobalamin (OH-B12) and other B-vitamins via liquid chromatography-mass spectrometry (LC-MS).

CeMbio isolates from -80 °C stocks were streaked on LB plates and incubated ON at 25°C. Per isolate, three single colonies were individually incubated in 4 ml LB using 48-well 6 ml polypropylene storage plates for 42 hours at 20°C on a circular vibrating shaker at 350 rpm. Bacterial inocula were transferred and centrifuged for 15 min at 3500 rpm. The supernatant was discarded, and the pellet was resuspended in 1 ml PBS and collected in a 1.5 ml Eppendorf tube before centrifugation for 5 min at 8000 rpm. The pellets’ wet weight was determined before storage at -20°C.

Cell pellets were resuspended in 750 µl methanol and transferred into 2 ml screwcap tups containing 1.0-1.2 mm diameter and 1.8-2.0 mm diameter SiLiBeads (Sigmund Lindner GmbH). Cells were disrupted by 2 cycles of bead beating for 30 sec at 7 m sec^-1^ divided by a 30-second gap. Samples were then centrifuged (13.000 rpm for 2 min at 4°C), and the supernatants transferred to new tubes. The pellets were extracted via bead beading again with 750 µl of freshly made extraction solution (2:2:1 acetonitrile: methanol: Milli-Q water). The supernatants were combined and evaporated to dryness in an Eppendorf Concentrator (SpeedVac) at 30°C for approximately 6 h. The obtained aliquots were stored at -80 °C until processing. For analysis, samples were resuspended with 100 µl of buffer A (0.1% formic acid in H_2_O). All samples were vortexed and centrifuged at 10.000 x g for 3 min at 4°C before mass spectrometry analysis. Samples were processed under natural light, which may impact measurement outcomes of the light sensitive vitamin B12.

Samples were analyzed on a Vanquish Horizon UHPLC system (Thermo Fisher Scientific) equipped with a binary pump, temperature-controlled autosampler (10°C), and column compartment (50°C) coupled to a QExactive Plus Orbitrap mass spectrometer (ThermoFischer Scientific) with a heated electrospray ionization probe (HESI-II). Chromatographic separation was achieved using a Accurcore ™ Vanquish C18+, 80 Å pore size, 1.5 μm particle size, 100 mm x 2.1 mm ID (27101-102130, Thermo-Scientific) column and mobile phases composed of 0.1% formic acid in H_2_O (A) and 0.1% formic acid in MeCN (B). The flow rate and injection volume were 0.4 ml/min and 5 μl, respectively. The gradient used was 0–10 min, 4–25% B; 10–11 min, 25-99% B; 11–12.5 min, 100% B; 12.5-13 min 100-4% B followed by a 2 min re-equilibration at 4% B.

Full-scan acquisition was performed from m/z 110-1000 in positive mode with an inclusion list of all forms of vitamin B12 and other B-vitamins. The mass resolution setting was 140.000, with an automatic gain control (AGC) target of 5 × 10^5^ and a maximum injection time of 50 ms per scan. The spray voltage of the source was +3.5 kV, with a capillary temperature of 300°C. The sheath and auxiliary gas were set at 50 and 10 (arbitrary units). The probe heater temperature was set at 450°C, and the S-Lens RF level was set to 60.

For quantification, we combined equal portions of each sample to obtain a quality control (QC) pool that was then used as an intra-run QC injection and the base for calibration curves, adjusted for fold-dilution. Calibration curves were performed in triplicate injections of 25, 50, 250, and 1000 fmol on an analytical column for all forms of B12. Calibration curves were used for quantification if R^2^ were all >0.99. Limits of detection (LOD) and limit of quantification (LOQ) were calculated as 3x and 10x the standard deviation of the lowest point of the matrix matched calibration curve.

### Vitamin B12 supplementation assay

Fertility assays followed by lifespan assays were conducted with non-B12-producing, single CeMbio isolates (*E. cloacae* CEnt1 and *A. guillouiae* MYb10) and compared to *E. coli* OP50 and B12-producing *P. lurida* MYb11 and *O. vermis* MYb71. We adapted a protocol from Bito et al. (2017)^94^ to generate B12-deficient worms. In this, worms were cultivated in V-medium with *E. coli* OP50 (final OD_600_ =1) for two generations at 20°C and 300 rpm.

Before each experiment, treatment V-medium was supplemented with methylcobalamin (MeCbl; 100 µg/l, 74.4 nM) (Roth, #13422-55-4). Worm age synchronization was achieved by bleaching gravid F2 hermaphrodites with an alkaline hypochlorite solution, as described above. Sterile, synchronized L1 larvae were directly transferred to single wells of 24-well plates each containing 1 ml V-medium inoculated with the respective CeMbio communities (final OD_600_ =0.1). Live adults were transferred daily to fresh plates seeded with the respective CeMbio strains (final OD_600_ =0.1) until the end of the egg-laying period, with offspring counts survival and scored daily until day of death. Experiments were conducted for one biological replicate consisting of separately cultured bacteria. Within each replicate, 22 technical replicates were cultured in parallel treatment plates to account for within-replicate variation.

### Vitamin B12 vs. non-B12 community assay

Nine members of the CeMbio collection were used to construct two communities: B12-consumers (B12C), consisting of *Ochrobactrum vermis* (MYb071), *Pseudomonas lurida* (MYb11), *Pseudomonas berkeleyensis* (MSPm1), and *Comamonas piscis* (BigB0172); and non-B12-consumers (NBC), comprising *Stenotrophomonas indicatrix* (JUb19), *Acinetobacter guillouiae* (MYb10), *Lelliottia amnigena* (JUb66), and *Enterobacter hormaechei* (Cent1). *Escherichia coli* OP50 was used for worm maintenance and adaptation to viscous medium.

To initiate cultures, 48-well plates containing 4 ml LB were inoculated with 4 µl of frozen stocks, sealed, and incubated at 20°C (300 rpm) for 42 hours. Post-incubation, communities were assembled by pooling pre-determined volumes of each strain^60^. The bacteria were washed three times by centrifugation (4000 rpm, 15 min) and resuspended in S-buffer. Finally, the optical density (OD_600_) was adjusted to 10.

To pre-adapt *C. elegans* to viscous medium, worms were cultured for two generations (7 days) in 6-well plates with 600 µl of OD_600_= 10 *E. coli* OP50.

Worms were bleached to synchronize the population, and approximately 100 eggs per well were transferred to 6-well plates containing 6 ml viscous medium and 600 µl of either B12C or NBC communities (OD_600_ = 10). After developing for two days to the L4 stage, 22 L4 worms per replicate (N=132 total) were transferred into individual 24-well plate wells containing 1 ml medium and 10 µl bacterial culture (final OD_600_=0.1). From day 1, worms were transferred daily to fresh 24-well plates to monitor survival and offspring counts.

## Results

To assess how BALO presence influences both the microbiota and the host, we monitored microbiota dynamics in *C. elegans* populations co-cultivated with defined microbiotas of varying diversity following a single pulse of either *Bdellovibrio* MYbb2, MYbb4, or a sterile buffer. This was followed by assays assessing host life-history traits.

### Broad prey-range BALO presence affects community composition and extends median host lifespan

We first assessed the effect of BALOs on microbiome diversity over time. *Bdellovibrio* DNA (via species-specific PCR or qPCR) or ASVs (via 16S amplicon sequencing) were consistently detected in BALO-inoculated worm and medium samples at low abundance, confirming predator persistence.

Interestingly, across all experiments, BALOs had no measurable effect on community composition in medium samples (Fig. S1).

In worms, CeMbio richness did not differ substantially between the two BALO treatments, but was significantly higher in the BALO MYbb2 treatment at days 3 and 21 (Fig. 1A).

**Fig. 1:**
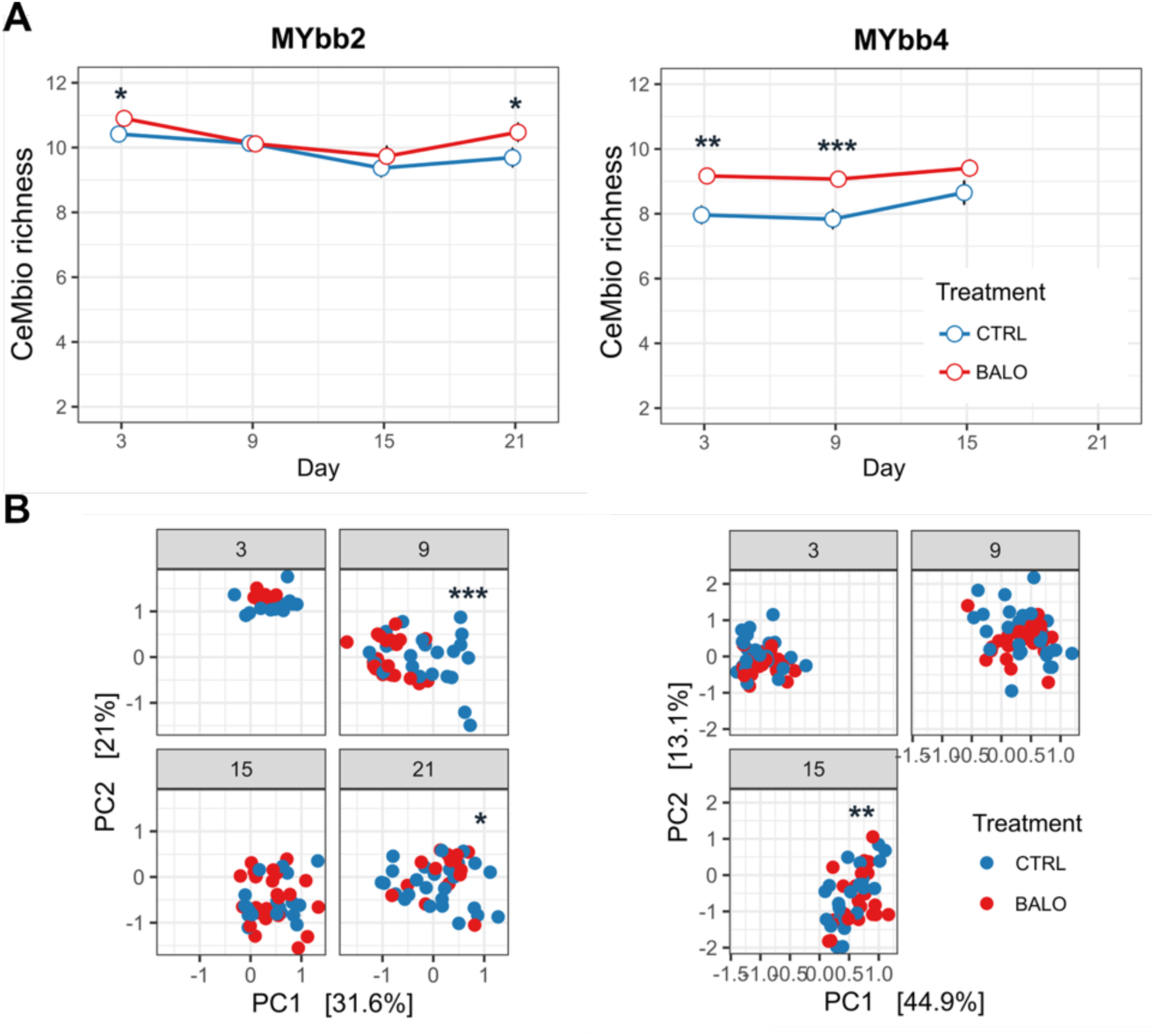
Diversity of worm-associated microbiomes in the longitudinal diversity experiments. (A) Alpha-diversity, shown as CeMbio richness, for samples from experiments with either the broad host-range BALO MYbb2 or the narrow host-range BALO MYbb4. Data averaged from all 6 biological replicates are shown. (B) Beta-diversity, shown as Aitchison distance, for samples from experiments with either MYbb2 or MYbb4. Each dot corresponds to one of the six biological replicates (each based on six technical replicates of 5 worms per sample). Asterisks indicate significant differences, calculated using (A) a generalized linear mixed model (GLMM) with a Generalized Poisson distribution (log link), with pairwise comparisons between treatments at each time point performed using estimated marginal means (EMMs), or (B) PERMANOVA, using technical replicates (i.e., individual worm samples) as a strata term to account for worms originating from the same biological replicate (i.e., flask).

The presence of *B. krueschi* MYbb4, which exclusively preys on *Ochrobactrum,* led to a similar pattern with significantly higher CeMbio richness on days 3 and 9.

Predator presence also modestly influenced beta diversity. Using Aitchison distance to account for the compositional nature of the data revealed a significant treatment effect on day 9, day 21 (MYbb2), and day 15 (MYbb4), respectively (Fig. 1B).

We next assessed whether the presence of BALOs (MYbb2 and MYbb4) translated into differences in host life-history traits. For this, three replicate populations per treatment were randomly selected for fitness assays, including lifespan and fertility (Fig. 2). Surprisingly, MYbb2 treatment significantly extended median lifespan compared to controls (where the mean lifespan was surprinsingly low: 10 days vs. 5 days; Cox mixed-effects model, p = 0.022), while mean offspring counts were higher but not significant. In contrast, MYbb4-treated worms showed no changes in lifespan or fertility, despite exhibiting increased microbiome diversity relative to controls. This suggests that richness per se does not explain the lifespan effect, but rather specific differences in community composition.

**Fig. 2:**
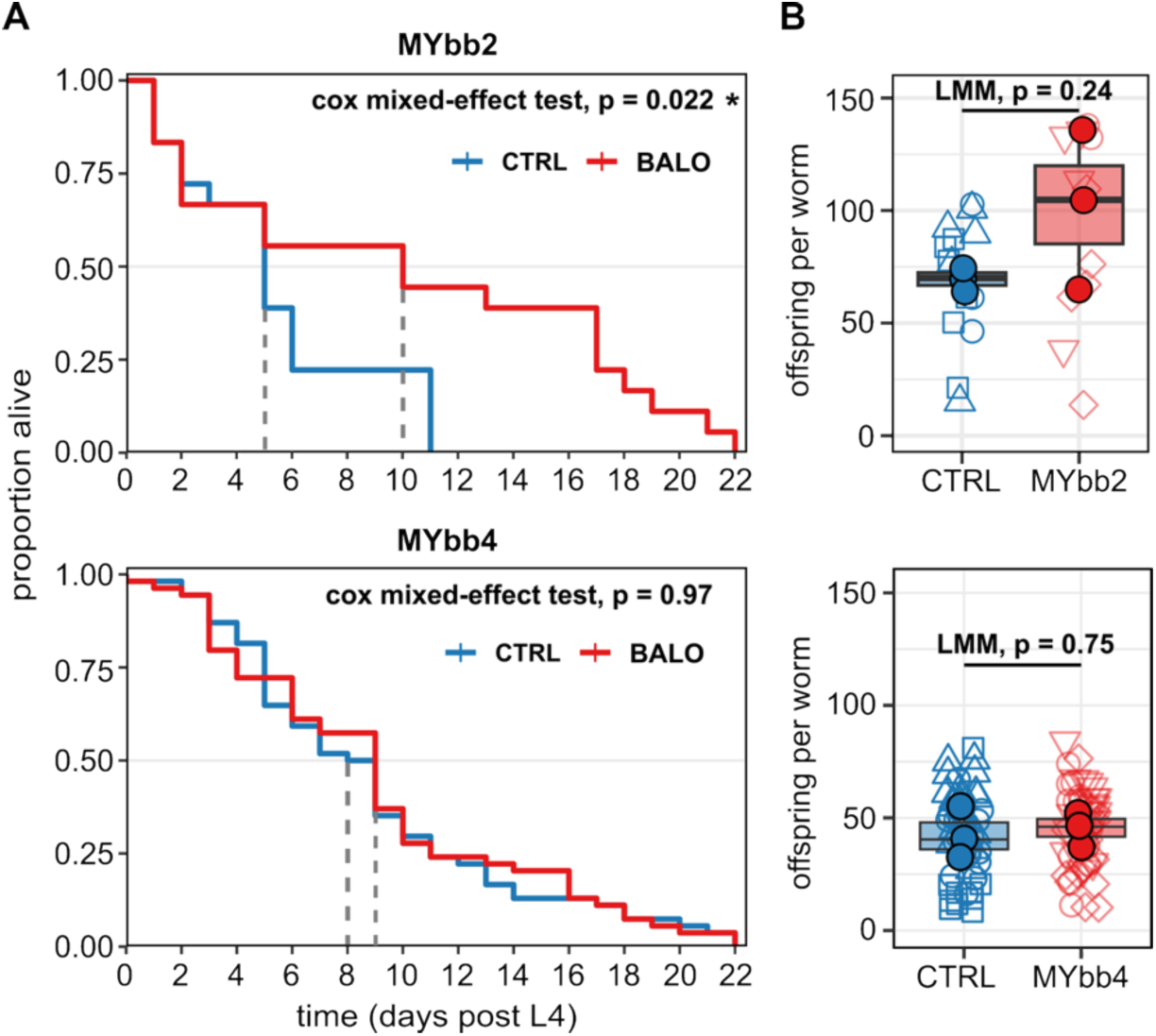
Host fitness outcomes following exposure to different BALO strains. Lifespan (A) and brood size (B) of worms exposed to the broad host-range BALO MYbb2 and the narrow host-range BALO MYbb4. A: Kaplan-Meier curves show median lifespan in worms compared to controls (Cox mixed-effects model). B: Mean offspring per worm across treatments, based on three biological replicates (n = 3). Unfilled symbols represent individual worms (technical replicates; 6 or 22 per biological replicate. Boxplots show replicate means. Groups were compared using a linear mixed effects model with biological replicate as random factor.

To further investigate the impact of predator exposure on the microbiome, we examined the community composition of worms from the three replicate flasks used for life-history analyses. Most notably, MYb71 (*Ochrobactrum*) increased in relative abundance in MYbb2-treated worms compared to both controls and MYbb4-treated worms (Fig. 3A, B; Fig. S2). Analysis of rank abundance distributions during the last sampling time point supported this shift in relative abundance and showed that MYb71 is the dominant member of the community in the MYbb2 but not the MYbb4 treatment (Fig. 3C and D). Among the differentially abundant taxa, MYb71 exhibited a large positive log-fold change in the MYbb2 treatment compared to control at day 21 (lfc = 0.76, ANCOMBC2). While this increase in MYb71 did not reach statistical significance (p.adj > 0.05), the magnitude of the shift was substantial, especially given the high relative abundance of MYb71. A post-hoc power analysis revealed an achieved power of only 44% (Cohen’s d = 1.94, Hedges’ g = 1.55, α = 0.05), indicating that more replicates would have been required to detect an significant effect of this magnitude. Given that MYb71 appeared to be the primary responder to the predator treatment, we subsequently focused our analysis on its metabolic potential to determine how its expansion might influence the community’s functional profile.

**Fig. 3:**
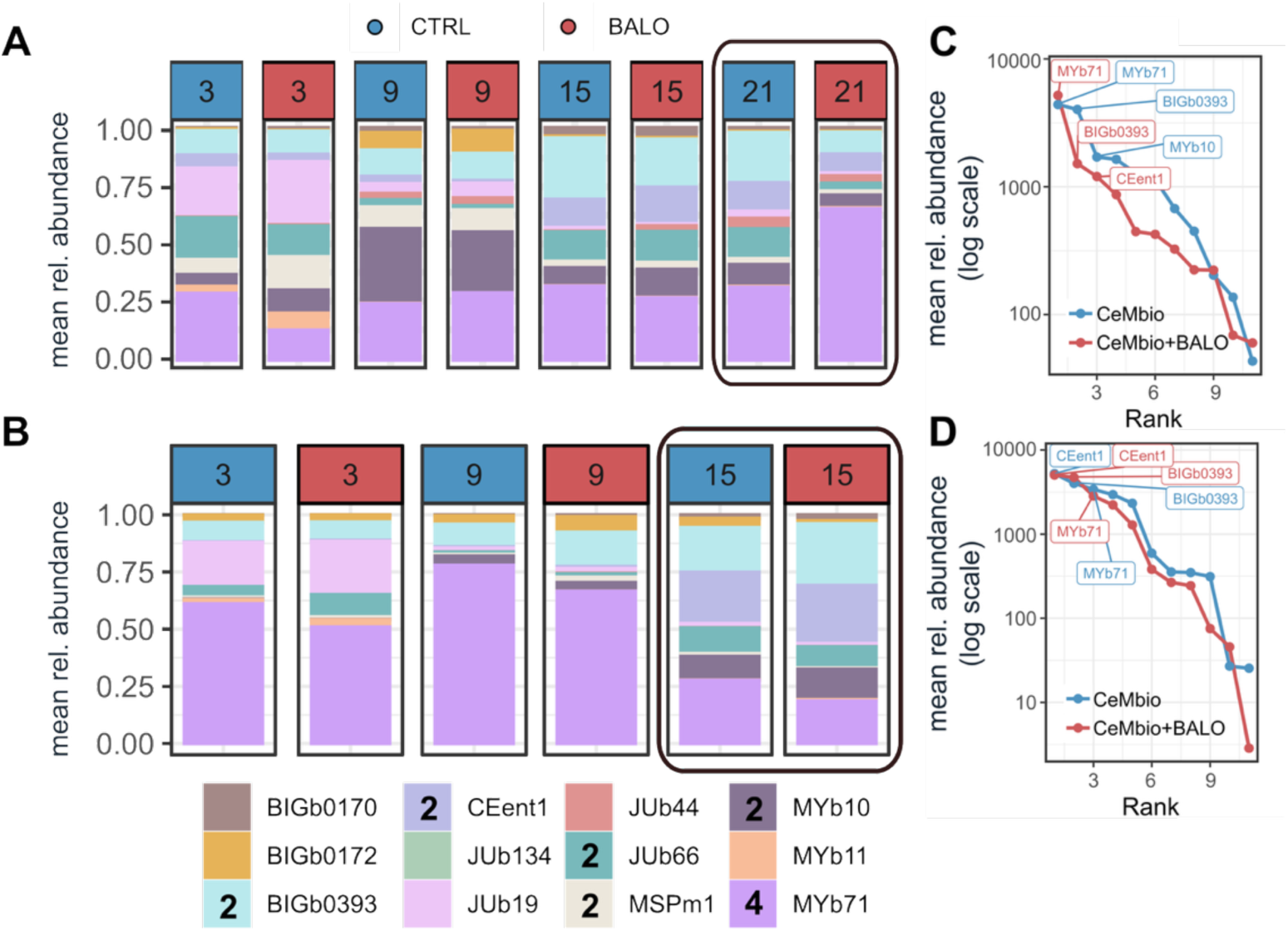
Temporal dynamics of *C. elegans* microbiome composition in response to BALO predation. Relative abundances of community members within the host microbiome were determined via 16S amplicon sequencing and adjusted for 16S rRNA gene copy numbers. One bar represents the mean of 6 technical replicate samples from the 3 biological replicates that were used for the fitness assay. (A) Microbiome composition in the presence of the broad host-range BALO MYbb2. (B) Microbiome composition in the presence of the narrow host-range BALO MYbb4. The final sampling timepoints are highlighted to emphasize long-term community shifts. Legend annotations indicate whether a specific CeMbio strain is susceptible to predation by MYbb2 (“2”) or MYbb4 (“4”). (C) and (D) Rank abundance plots corresponding to the highlighted last sampling day for the MYbb2 (C) and MYbb4 (D) experiment. The top three ranked taxa are labeled.

It should be noted that the observed pattern is consistent with the ecological interactions within the community: MYb71 is not preyed upon by MYbb2, allowing it to reach high abundance over time, whereas it is the only prey of MYbb4, which likely constrains its population size steadily (Fig. 3B). Notably, MYbb4 treatment, which suppresses MYb71 abundance through predation, did not result in lifespan differences, further supporting a potential role of this strain in mediating the observed phenotype.

We therefore hypothesized that MYb71 abundance, rather than *Bdellovibrio* presence per se, is a key factor underlying the observed lifespan differences. To explore potential mechanisms, we next considered metabolites produced by MYb71. Given that the complete genome sequence of MYb71 is available, biosynthetic pathways can be predicted^87^. We therefore surveyed the literature for microbial metabolites known to influence *C. elegans* lifespan.

Several metabolites have been implicated in lifespan regulation. For instance, altered bacterial folate synthesis and host one-carbon folate flux can modulate lifespan^95–97^. Tryptophan and microbiota-derived indoles can promote healthspan in an AHR-1-dependent manner, but their effects are metabolite- and context-dependent^13,98^. Additionally, serine, glycine, and methionine metabolism feed into one-carbon metabolism and methylation capacity, and methionine-restriction-like states are often associated with lifespan extension and altered reproductive output^96,97^. However, these candidates are unlikely to explain our observations. Folate and tryptophan biosynthesis pathways are predicted to be present in all CeMbio strains, and indole production is restricted to *Chryseobacterium* JUb44, which is not abundant in our dataset (Fig. S3). Similarly, methionine metabolism is broadly conserved across the community, making it an unlikely driver. Bacterial siderophores produced by BALOs and CeMbio members can influence host iron homeostasis and mitochondrial physiology^99^, but given their low abundance, this mechanism is unlikely to account for the observed phenotype.

In contrast, B vitamins are strong candidates, as bacterially supplied B-vitamins can directly shape host metabolism, development, and other life-history traits in *C. elegans*^95^. Vitamins B2 and B6, for example, support metabolic homeostasis and postembryonic development, linking bacterial micronutrient supply to host physiology^100,101^. Vitamin B12 (cobalamin), in particular, is critical for mitochondrial propionate breakdown and one-carbon metabolism in *C. elegans*, and altered B12 availability has been shown to affect fertility, development, stress physiology, and lifespan-related phenotypes depending on context^102–104^. We therefore quantified relevant B-vitamin production by CeMbio strains using targeted metabolomics (Fig. 4A, Fig. S6). MYb71 was found to produce both vitamin B6 and B12. While several CeMbio strains can produce B-vitamins, B12 biosynthesis is restricted to only four members of the community, with MYb71 being by far the most abundant in the worm microbiome (Fig. 3 A and B). When estimating community-level vitamin availability based on relative abundances at the final sampling time point, we found that only vitamin B12 levels were increased in the MYbb2 treatment compared to controls (approximately two-fold; Fig. 4B).

**Fig. 4:**
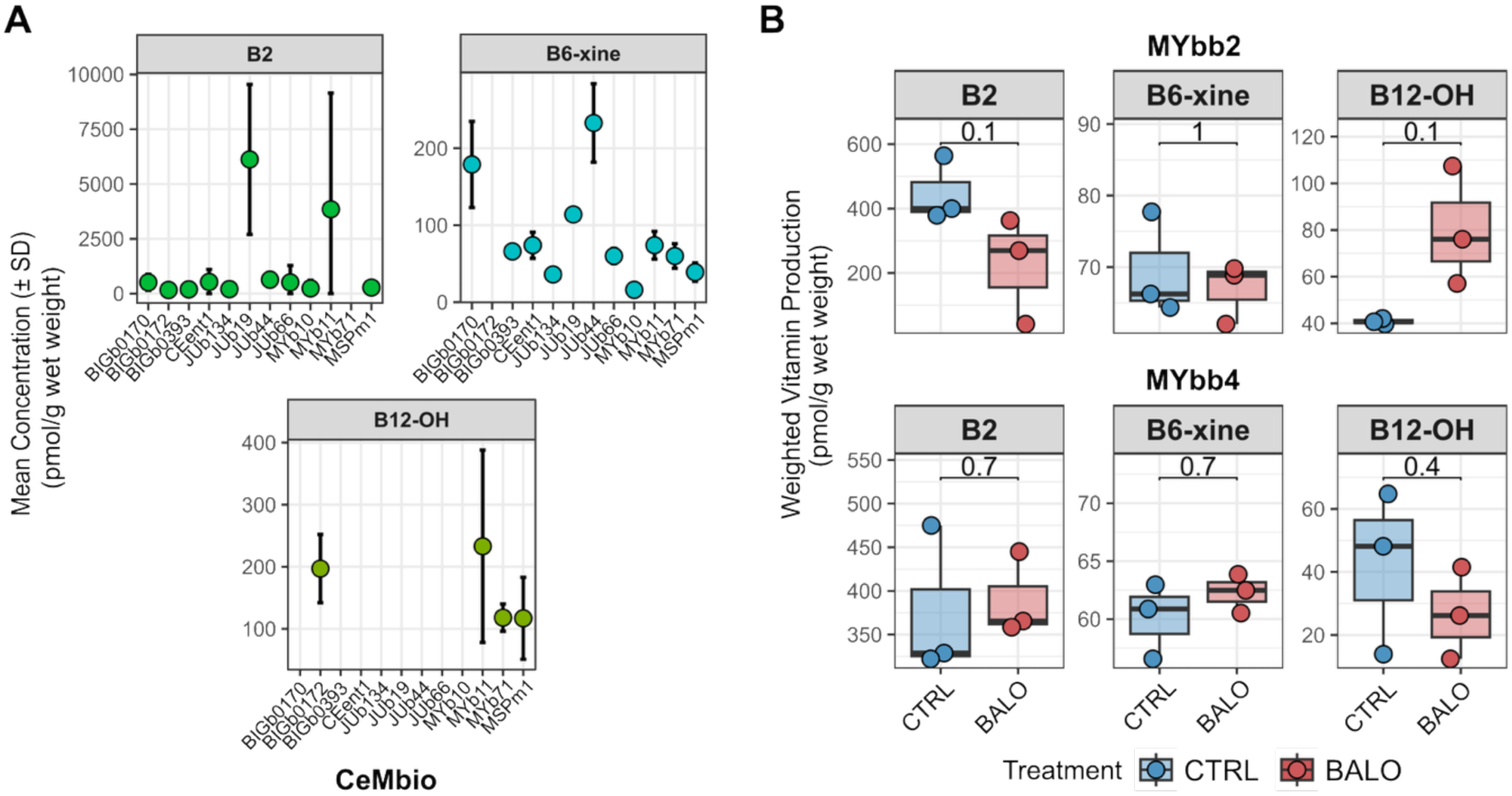
Vitamin production by CeMbio strains (A) and relative abundance of vitamins in CeMbio communities (B). A: Concentration (pmol of molecule extracted per gram of wet weight) of each vitamin per bacterial species. Only samples were at least 2 out of 3 biological replicates were above the limit of quantification are shown. Samples that fell below limit of quantification were removed. B: Weighted vitamin production of worm microbiomes based on the relative abundances of CeMbio strains during the last sampling time point and the mean vitamin production per CeMbio strain. A Wilcoxon test was used to calculate the statistics.

These results suggest that elevated vitamin B12 availability may contribute to the extended lifespan observed in the MYbb2 treatment. In *C. elegans*, vitamin B12 is required for efficient canonical propionate breakdown; when B12 is limiting, propionate accumulates and induces *acdh-1* and other propionate-shunt genes, consistent with elevated metabolic and mitochondrial stress^103,105,106^. Although mild mitochondrial stress can have hormetic effects, chronic stress is generally detrimental to fitness^107,108^.

Although BALOs themselves may contribute to propionate metabolism, their low abundance suggests a limited direct effect. In contrast, CeMbio members capable of producing propionate could impose a metabolic burden on the host, increasing the need for B12-mediated detoxification. Based on genomic predictions, we identified CeMbio strains capable of propionate fermentation (Fig. S3) and calculated the ratio of B12 producers to propionate fermenters using relative abundances at the final time point (Fig. 5). It should be noted that this categorical assignment assumes that metabolic potential correlates with strain abundance and does not account for actual metabolite flux under differing environmental conditions. Notably, this ratio was highest in the MYbb2 treatment, indicating a community shift toward surplus B12 supply relative to potential propionate-mediated metabolic demand, consistent with the observed extensions in host median lifespan and elevated fertility.

**Figure 5:**
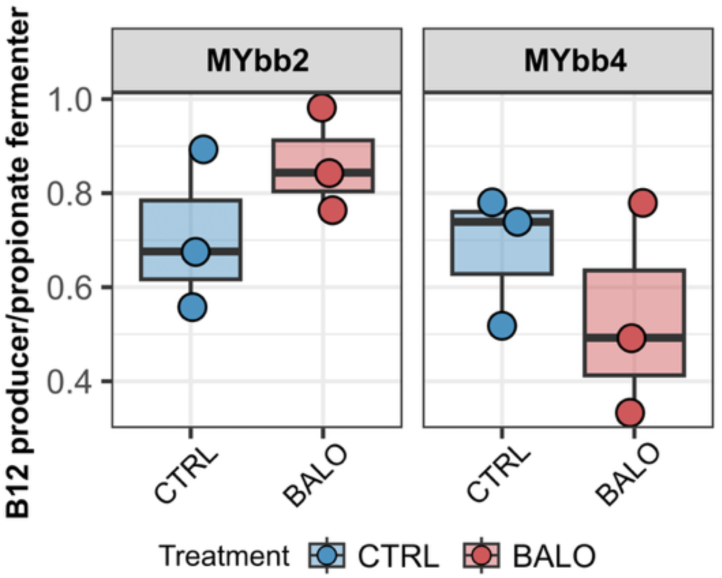
Ratio of the summed relative abundance of vitamin B12 producers to propionate fermenters in worm microbiomes at the final sampling time point (Day 21 for MYbb2, Day 15 for MYbb4). Fermentation and B12 biosynthesis capacities were assigned based on genomic predictions for each CeMbio strain. Statistical significance was determined using [Wilcoxon rank-sum test], with significance levels indicated as follows: * p < 0.05, ** p < 0.01.

In contrast, the MYbb4 treatment showed a lower B12-to-propionate ratio at the final time point, yet did not result in a shorter lifespan relative to the CeMbio control. This apparent discrepancy may be explained by the temporal dynamics of MYb71 abundance: while MYb71 was suppressed by MYbb4 predation at later time points, it dominated the community at earlier stages of the experiment (Fig. 3B). Given that vitamin B12 is maternally transmitted in *C. elegans*^109^, the B12 acquired by earlier generations may have been sufficient to sustain host fitness in subsequent generations, even as community B12 production capacity declined. In other words, the historically high B12 availability may buffer the worms against the later reduction in B12 producers.

This interpretation is further supported by the contrasting temporal dynamics observed in the MYbb2 treatment, where MYb71 abundance increased steadily over time (Fig. 3A), providing a growing source of B12 throughout the experiment. Together, these observations suggest that it is not only the instantaneous community composition, but also its trajectory over time, that shapes host life-history outcomes.

Together, these observations support a model in which the balance between B12 production and propionate metabolism, integrated over the course of microbiota succession, contributes to host life-history outcomes.

### Bacterial vitamin B12 production affects host life-history traits

We next screened all 12 CeMbio strains using the dietary sensor strain *C. elegans acdh-1*p::GFP^65^. In this reporter, expression of *acdh-1*, which encodes an acyl-CoA dehydrogenase involved in the mitochondrial propionate shunt, is downregulated in the presence of B12, resulting in the absence of a GFP signal^21,65,93^. Thus, this assay indicates whether worms obtain vitamin B12 from a given bacterial strain.

Consistent with predictions, worms fed with *P. lurida* (MYb11), *O. vermis* (MYb71) and *C. piscis* (BIGb0172), and *P. berkeleyensis* (MsPM1) showed no GFP signal, confirming microbial B12 production (Fig. 6).

**Figure 6:**
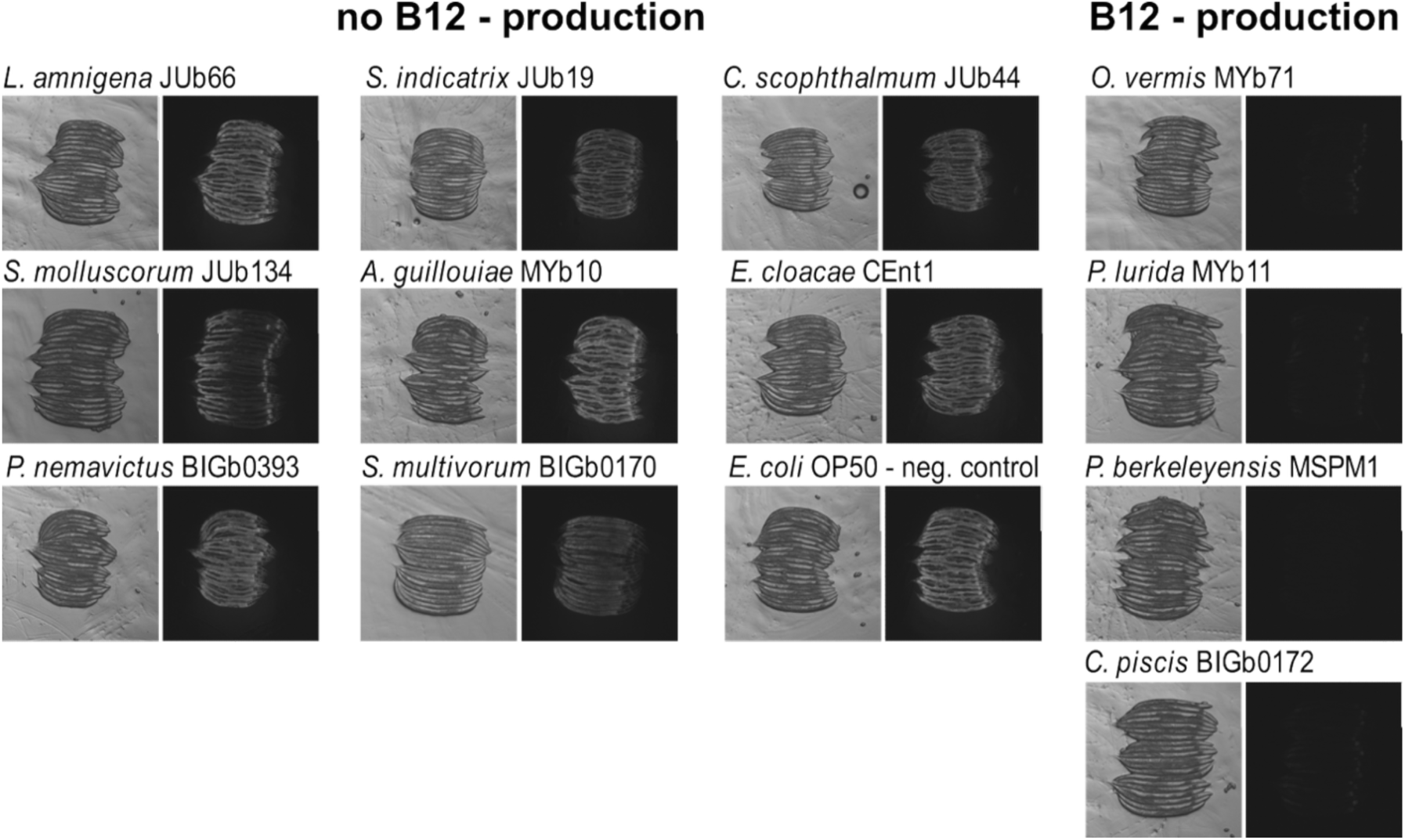
GFP signal of transgenic C. elegansreporter strain acdh-1p::gfp in response to bacterial vitamin B12 production. (A) Representative images of worms were exposed to either E. coli OP50 or one of the 12 CeMbio strains. Worms were imaged as young adults in groups of approximately 20, arranged with heads oriented to the right. Each panel shows the corresponding transmitted light image (left) and GFP fluorescence (right).

Since increased abundance of B12-producing bacteria correlated with extended lifespan, we next tested whether exogenous methylcobalamin (MeCbl), a biologically active form of vitamin B12, could influence host life-history traits across different bacterial diets. Lifespan and fertility assays were performed using worms grown on individual bacterial isolates with or without MeCbl supplementation (Fig. 7).

**Fig. 7:**
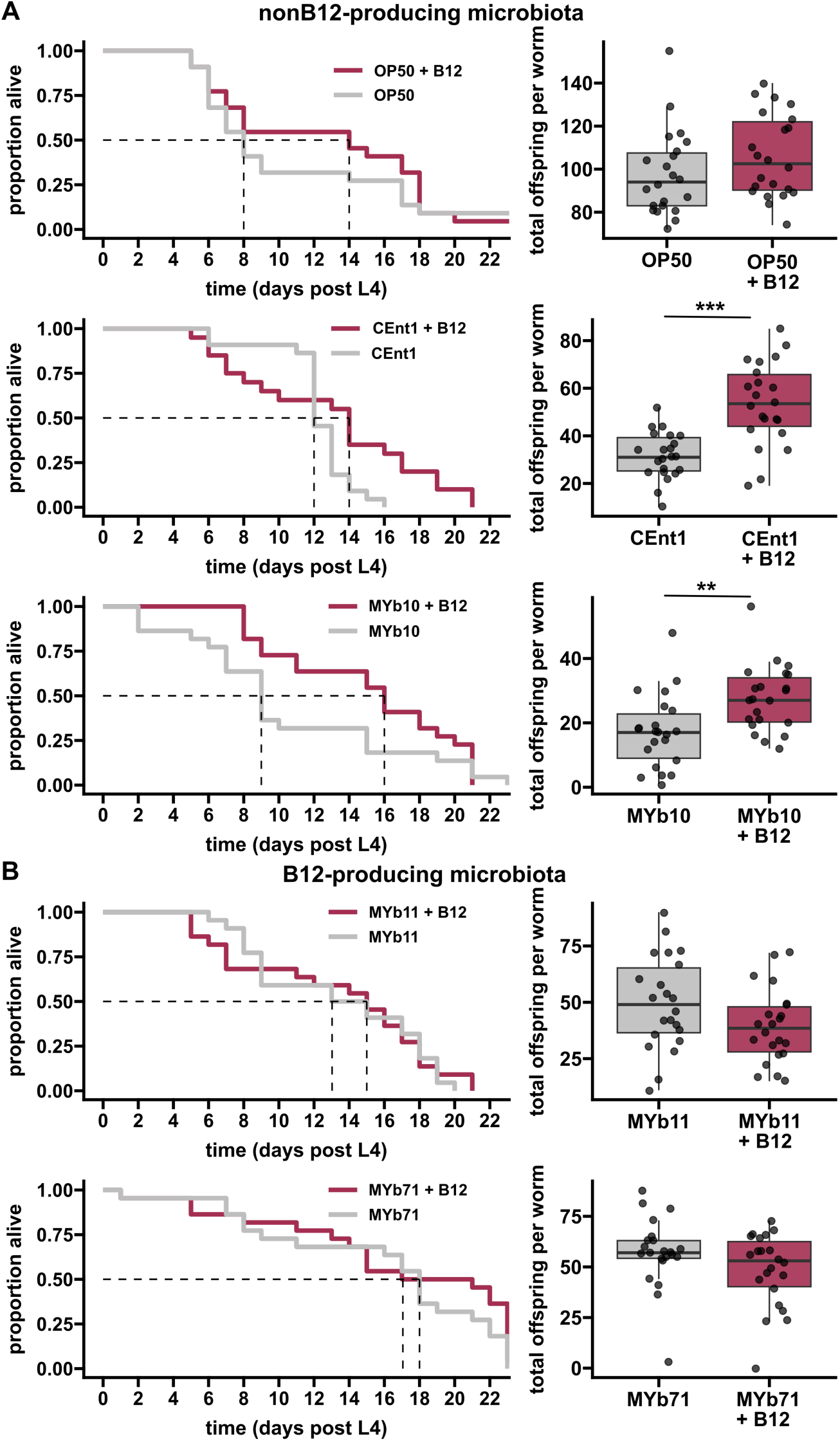
Effects of exogenous vitamin B12 supplementation on *C. elegans* lifespan and brood size. Each condition was tested with 22 worms. (A) Supplementation with exogenous B12 significantly increases both lifespan and brood size in worms fed non-B12-producing bacteria. (B) B12 supplementation has no effect on lifespan or brood size in worms fed B12-producing bacteria, indicating that B12 availability is already saturating under these conditions. All experiments were conducted in viscous medium. Brood size data were analyzed using the Wilcoxon test.

In non-B12-producing diets (*E. coli* OP50, *E. cloacae* CEent1, and *A. guillouiae* MYb10), MeCbl supplementation increased both median lifespan and offspring production (Fig. 6A–C). In contrast, supplementation had no effect on lifespan or fertility in worms fed B12-producing strains (*P. lurida* MYb11 and *O. vermis* MYb71; Fig. 7D–E), suggesting that endogenous B12 production may already meet or exceed host requirements. To test if B12 production is universally sufficient to extend lifespan, we designed synthetic communities composed of either de novo producers or salvagers. Interestingly, we found no lifespan differences in this simplified context, where both communities supported high host longevity (Fig. S4). However, worm fertility was slightly increased in the B12-producing community. This suggests that B12 availability is not always a limiting factor. However, in the context of the more complex and ecologically dynamic longitudinal experiments, where baseline lifespan was unexpectedly low in the control treatment (see Fig. 2), the predator-induced enrichment of B12-producers appears to resolve a specific metabolic bottleneck.

To further investigate the robustness of the lifespan effect and its potential maternal transmission, we expanded our experimental system to a complex standardized microbiota of 43 bacterial strains (CeMbio43)^60^. In this setup, we again observed a significant enrichment of *Ochrobactrum* in the microbiome of parent worms following MYbb2 treatment without striking differences in microbiome richness (Fig. S5 A and B).

We then performed a bleaching protocol to isolate embryos from these populations, effectively removing the parent-associated microbiota and ensuring that any phenotypic effects in the offspring would be driven by maternal provisioning or the re-establishment of the microbiota. Interestingly, the resulting F1 generation still exhibited an increased lifespan compared to controls (Fig. S5 C).

Given that vitamin B12 is known to be maternally provisioned from *C. elegans* mothers to their embryos, these results suggest that the MYbb2-induced dominance of B12-producing bacteria in the parental generation is sufficient to improve life-history traits in the subsequent generation.

## Discussion

While prior work has highlighted BALOs as potential biocontrol agents^48^ or drivers of turnover in free-living bacterial populations^110^, their impacts on the diversity, composition, and function of host-associated microbiotas remains underexplored. Our findings experimentally demonstrate that bacterial predators can restructure host-associated microbiotas and modulate host fitness through direct and indirect interactions with beneficial bacteria. By comparing the impact of two BALO species that differ in prey range, the broad-spectrum predator *B. tiberii* MYbb2 and the narrow-spectrum *B. krueschi* MYbb4, on a defined *C. elegans*-associated microbiota (CeMbio), we demonstrate that predator-induced structural and functional shifts are dependent on prey specificity and initial microbiota composition. Using the well-characterized *C. elegans* model system allowed us to identify the molecular mechanism underlying these effects: the production of vitamin B-12.

The narrow-range predator MYbb4, which targets *Ochrobactrum* MYb71, significantly increased community alpha diversity. This suggests that MYbb4 acts as a top-down regulator, potentially increasing diversity by suppressing the dominant *Ochrobactrum* strain and liberating niche space for less abundant taxa. Notably, *Ochrobactrum* is a ubiquitous and robust colonizer of the *C. elegans* gut, frequently identified as a member of the worm’s “core microbiome”^53,57,59^. Despite these structural shifts, MYbb4-mediated predation did not measurably alter the host life-history traits assayed, indicating that increased microbiome diversity does not inherently translate to altered host fitness. In contrast, the broad-range predator MYbb2 induced modest diversity changes but significantly extended *C. elegans* median lifespan. Intriguingly, MYbb2 presence correlated with an enrichment of *Ochrobactrum* MYb71. Given that *Ochrobactrum* is established as a persistent commensal that modulates host insulin signaling, enhances growth, and buffers against environmental stress^17,20,59,87^, our results suggest that MYbb2 may indirectly improve host fitness by selectively favoring functionally beneficial “keystone” taxa.

A primary mechanism for this benefit appears to be the provisioning of vitamin B12 (cobalamin), for which *Ochrobactrum* is one of only four producers in the CeMbio community. Because vitamin B12 is an essential cofactor for DNA synthesis and mitochondrial function, and specifically regulates the propionate degradation pathway in *C. elegans*^65,93,105^, its availability has direct implications for development and longevity. While targeted supplementation with methylcobalamin (MeCbl) consistently improved lifespan and fertility across multiple bacterial backgrounds (OP50, CEnt1, MYb10), the relationship between B12 and host health is clearly non-linear and context-dependent. Our data suggest that while B12 broadly supports fertility, its impact on lifespan is modulated by the metabolic environment, particularly the burden of microbiota-produced propionate. In *C. elegans*, insufficient B12 leads to the accumulation of toxic propionate via the shunt pathway^102^. The observed shift in the B12:propionate ratio in MYbb2-treated communities suggests that predation helps maintain B12 levels sufficient to detoxify metabolic byproducts, thereby preventing the lifespan reduction associated with propionate-induced stress. This explains why simplified communities of B12-producers alone were sufficient to increase offspring production but did not further extend lifespan: the already high lifespan in non-B12 communities suggests that propionate accumulation was not a problem in these contexts. Beyond B12, our LC-MS analysis reveals a pattern of metabolic niche partitioning: B12 producers rarely co-produce other essential vitamins, such as biotin (vitamin B7, Fig. S6). This implies that host fitness is supported by a distributed metabolic network rather than single-strain effects^111,112^. Coordinated production of different vitamins, such as B12 and B6, may drive the synergistic host responses previously observed in multispecies bacterial communities^20^. Furthermore, predator-induced cell lysis may facilitate this cooperation by releasing intracellular vitamers into the extracellular environment, effectively “feeding” cross-feeding networks, a phenomenon previously documented in phage-bacteria interactions involving B12^113^.

Finally, we ruled out dietary restriction (DR) as a confounding mechanism. While predation inherently reduces bacterial biomass, which could theoretically trigger DR-mediated longevity^114,115^, this is not supported by our comparative data. If biomass reduction alone were responsible, both MYbb2 and MYbb4 would be expected to extend lifespan. The fact that only MYbb2 conferred a longevity benefit reinforces a model predicated on selective community restructuring and the enrichment of specific metabolic functions.

## Conclusion and Outlook

In conclusion, BALOs are potent ecological regulators capable of enhancing host life-history traits through the targeted enrichment of beneficial microbial taxa. These effects are highly specific to the predator’s prey range and the existing microbiota structure. Moving forward, resolving the precise spatiotemporal dynamics of predator-prey-host interactions will be critical for harnessing BALOs as precision microbiome modulators. Our study provides a foundation for developing predator-based probiotic strategies aimed at restoring or enhancing the metabolic output of host-associated microbiotas.

## Acknowledgements

We thank the Schulenburg group for discussions and advice on this work. We thank Lisa Klameth, Barbara Pees, Yvonne Carstensen, Halil Furkan Tuna, Katharina Rathjen, and Laila Jacobsen for their valuable contributions to this work. For genomic sequencing, we further thank the Competence Centre for Genomic Analysis (CCGA) Kiel. We are grateful for funding from the German Research Foundation (Deutsche Forschungsgemeinschaft, DFG) within the DFG project JO 1786/1-1 (to JJ) and within the Collaborative Research Center (CRC) 1182 on the Origin and Function of Metaorganisms (project Z3 to ML and Mercator fellowship to BB). We gratefully acknowledge funding from the Max Planck Society (to CB and ML).

## Data availability

Sequences are publicly available at the SRA under accession number PRJNA1481537.

## References

1. Bosch, T. C. G. & McFall-Ngai, M. J. Metaorganisms as the new frontier. Zoology 114, 185–190 (2011).

2. McFall-Ngai, M. et al. Animals in a bacterial world, a new imperative for the life sciences. Proc. Natl. Acad. Sci. 110, 3229–3236 (2013).

3. Cabreiro, F. & Gems, D. Worms need microbes too: microbiota, health and aging in *Caenorhabditis elegans*. Embo Mol. Med. 5, 1300–1310 (2013).

4. Rowland, I. et al. Gut microbiota functions: metabolism of nutrients and other food components. Eur. J. Nutr. 57, 1–24 (2018).

5. Warnecke, F. et al. Metagenomic and functional analysis of hindgut microbiota of a wood-feeding higher termite. Nature 450, 560–565 (2007).

6. Bäckhed, F., Ley, R. E., Sonnenburg, J. L., Peterson, D. A. & Gordon, J. I. Host-bacterial mutualism in the human intestine. Science 307, 1915–1920 (2005).

7. Bosch, T. C. G. Rethinking the role of immunity: lessons from *Hydra*. Trends Immunol. 35, 495–502 (2014).

8. Jia, Y. et al. Gut microbiome modulates *Drosophila* aggression through octopamine signaling. Nat. Commun. 12, 2698 (2021).

9. Nagpal, J. & Cryan, J. F. Microbiota-brain interactions: Moving toward mechanisms in model organisms. Neuron 109, 3930–3953 (2021).

10. Ruff, W. E., Greiling, T. M. & Kriegel, M. A. Host-microbiota interactions in immune-mediated diseases. Nat. Rev. Microbiol. 18, 521–538 (2020).

11. Sommer, F. & Bäckhed, F. The gut microbiota--masters of host development and physiology. Nat. Rev. Microbiol. 11, 227–238 (2013).

12. Leitão-Gonçalves, R. et al. Commensal bacteria and essential amino acids control food choice behavior and reproduction. PLOS Biol. 15, e2000862 (2017).

13. Sonowal, R. et al. Indoles from commensal bacteria extend healthspan. Proc. Natl. Acad. Sci. U. S. A. 114, E7506–E7515 (2017).

14. Round, J. L. & Mazmanian, S. K. The gut microbiota shapes intestinal immune responses during health and disease. Nat. Rev. Immunol. 9, 313–323 (2009).

15. González, R. & Félix, M.-A. Naturally-associated bacteria modulate Orsay virus infection of *Caenorhabditis elegans*. PLOS Pathog. 20, e1011947 (2024).

16. King, K. C. et al. Rapid evolution of microbe-mediated protection against pathogens in a worm host. ISME J. 10, 1915–1924 (2016).

17. Obeng, N., Bansept, F., Sieber, M., Traulsen, A. & Schulenburg, H. Evolution of microbiota-host associations: The microbe’s perspective. Trends Microbiol. 29, 779–787 (2021).

18. Shapira, M. Host–microbiota interactions in *Caenorhabditis elegans* and their significance. Curr. Opin. Microbiol. 38, 142–147 (2017).

19. Vuong, H. E., Yano, J. M., Fung, T. C. & Hsiao, E. Y. The microbiome and host behavior. Annu. Rev. Neurosci. 40, 21–49 (2017).

20. Haçariz, O., Viau, C., Karimian, F. & Xia, J. The symbiotic relationship between *Caenorhabditis elegans* and members of its microbiome contributes to worm fitness and lifespan extension. BMC Genomics 22, 364 (2021).

21. Watson, E. et al. Interspecies systems biology uncovers metabolites affecting *C. elegans* gene expression and life history traits. Cell 156, 759–770 (2014).

22. Griem-Krey, H., Petersen, C., Hamerich, I. K. & Schulenburg, H. The intricate triangular interaction between protective microbe, pathogen and host determines fitness of the metaorganism. Proc. R. Soc. B Biol. Sci. 290, 20232193 (2023).

23. Kissoyan, K. A. B. et al. Natural *C. elegans* microbiota protects against infection via production of a cyclic lipopeptide of the viscosin group. Curr. Biol. 29, 1030–1037.e5 (2019).

24. Kissoyan, K. A. B. et al. Exploring effects of *C. elegans* protective natural microbiota on host physiology. Front. Cell. Infect. Microbiol. 12, 775728 (2022).

25. Pees, B. et al. The *C. elegans* proteome response to two protective *Pseudomonas* mutualists. 2023.03.22.533766 Preprint at 10.1101/2023.03.22.533766 (2024).

26. Peters, L. et al. Polyketide synthase-derived sphingolipids mediate microbiota protection against a bacterial pathogen in *C. elegans*. Nat. Commun. 16, 5151 (2025).

27. Sommer, F., Anderson, J. M., Bharti, R., Raes, J. & Rosenstiel, P. The resilience of the intestinal microbiota influences health and disease. Nat. Rev. Microbiol. 15, 630–638 (2017).

28. Nguyen, J., Lara-Gutiérrez, J. & Stocker, R. Environmental fluctuations and their effects on microbial communities, populations and individuals. FEMS Microbiol. Rev. 45, fuaa068 (2021).

29. Tropini, C. How the physical environment shapes the microbiota. mSystems 6, 10.1128/msystems.00675-21 (2021).

30. Sieber, M., Traulsen, A., Schulenburg, H. & Douglas, A. E. On the evolutionary origins of host–microbe associations. Proc. Natl. Acad. Sci. 118, e2016487118 (2021).

31. Taylor, M. & Vega, N. M. Host immunity alters community ecology and stability of the microbiome in a *Caenorhabditis elegans* model. mSystems 6, e00608–20 (2021).

32. Clemente, J. C., Ursell, L. K., Parfrey, L. W. & Knight, R. The impact of the gut microbiota on human health: an integrative view. Cell 148, 1258–1270 (2012).

33. Levy, M., Kolodziejczyk, A. A., Thaiss, C. A. & Elinav, E. Dysbiosis and the immune system. Nat. Rev. Immunol. 17, 219–232 (2017).

34. Wilde, J., Slack, E. & Foster, K. R. Host control of the microbiome: Mechanisms, evolution, and disease. Science 385, eadi3338 (2024).

35. Peixoto, R. S., Rosado, P. M., Leite, D. C. de A., Rosado, A. S. & Bourne, D. G. Beneficial Microorganisms for Corals (BMC): Proposed mechanisms for coral health and resilience. Front. Microbiol. 8, (2017).

36. Matsuoka, K. & Kanai, T. The gut microbiota and inflammatory bowel disease. Semin. Immunopathol. 37, 47–55 (2015).

37. Zheng, D., Liwinski, T. & Elinav, E. Interaction between microbiota and immunity in health and disease. Cell Res. 30, 492–506 (2020).

38. Pamer, E. G. Fecal microbiota transplantation: effectiveness, complexities, and lingering concerns. Mucosal Immunol. 7, 210–214 (2014).

39. Kristensen, N. B. et al. Alterations in fecal microbiota composition by probiotic supplementation in healthy adults: a systematic review of randomized controlled trials. Genome Med. 8, 52 (2016).

40. Walter, J., Maldonado-Gómez, M. X. & Martínez, I. To engraft or not to engraft: an ecological framework for gut microbiome modulation with live microbes. Curr. Opin. Biotechnol. 49, 129–139 (2018).

41. Dosoky, N. S., May-Zhang, L. S. & Davies, S. S. Engineering the gut microbiota to treat chronic diseases. Appl. Microbiol. Biotechnol. 104, 7657–7671 (2020).

42. Sanders, M. E. Probiotics: considerations for human health. Nutr. Rev. 61, 91–99 (2003).

43. Vyas, U. & Ranganathan, N. Probiotics, prebiotics, and synbiotics: gut and beyond. Gastroenterol. Res. Pract. 2012, 872716 (2012).

44. Jurkevitch, E. The genus *Bdellovibrio*. in The Prokaryotes (eds Dworkin, M., Falkow, S., Rosenberg, E., Schleifer, K.-H. & Stackebrandt, E.) 12–30 (Springer New York, New York, NY, 2006). doi:10.1007/0-387-30747-8_2.

45. Welsh, R. M. et al. Alien vs. predator: bacterial challenge alters coral microbiomes unless controlled by *Halobacteriovorax* predators. PeerJ 5, e3315 (2017).

46. Winter, C., Bouvier, T., Weinbauer, M. G. & Thingstad, T. F. Trade-offs between competition and defense specialists among unicellular planktonic organisms: the ‘killing the winner’ hypothesis revisited. Microbiol. Mol. Biol. Rev. 74, 42–57 (2010).

47. Bonfiglio, G. et al. Insight into the possible use of the predator *Bdellovibrio bacteriovorus* as a probiotic. Nutrients 12, 2252 (2020).

48. Bratanis, E., Andersson, T., Lood, R. & Bukowska-Faniband, E. Biotechnological potential of *Bdellovibrio* and like organisms and their secreted enzymes. Front. Microbiol. 11, (2020).

49. Cavallo, F. M., Jordana, L., Friedrich, A. W., Glasner, C. & Dijl, J. M. van. *Bdellovibrio bacteriovorus*: a potential ‘living antibiotic’ to control bacterial pathogens. Crit. Rev. Microbiol. 0, 1–17 (2021).

50. Pérez, J., Contreras-Moreno, F. J., Marcos-Torres, F. J., Moraleda-Muñoz, A. & Muñoz-Dorado, J. The antibiotic crisis: How bacterial predators can help. Comput. Struct. Biotechnol. J. 18, 2547–2555 (2020).

51. Pasternak, Z. et al. In and out: an analysis of epibiotic vs periplasmic bacterial predators. ISME J. 8, 625–635 (2014).

52. Maher, R. L. et al. Comparative analysis of novel Pseudobdellovibrionaceae genera and species yields insights into the genomics and evolution of bacterial predation mode. 2025.02.19.638989 Preprint at 10.1101/2025.02.19.638989 (2025).

53. Johnke, J., Dirksen, P. & Schulenburg, H. Community assembly of the native *C*. *elegans* microbiome is influenced by time, substrate and individual bacterial taxa. Environ. Microbiol. 22, 1265–1279 (2020).

54. Hoffmann, A., Müller, T., Fingerle, V. & Noll, M. Presence of human pathogens of the *Borrelia burgdorferi* sensu lato complex shifts the sequence read abundances of tick microbiomes in two German locations. Microorganisms 9, 1814 (2021).

55. Iebba, V. et al. Higher prevalence and abundance of *Bdellovibrio bacteriovorus* in the human gut of healthy subjects. Plos One 8, (2013).

56. Johnke, J., Fraune, S., Bosch, T. C. G., Hentschel, U. & Schulenburg, H. *Bdellovibrio* and like organisms are predictors of microbiome diversity in distinct host groups. Microb. Ecol. 79, 252–257 (2020).

57. Dirksen, P. et al. The native microbiome of the nematode *Caenorhabditis elegans*: gateway to a new host-microbiome model. BMC Biol. 14, 38 (2016).

58. Johnke, J., et al. *Caenorhabditis* nematodes influence microbiome and metabolome characteristics of their natural apple substrates over time. mSystems 0, e01533–24 (2025).

59. Dirksen, P. et al. CeMbio - The *Caenorhabditis elegans* microbiome resource. G3 GenesGenomesGenetics 10, 3025–3039 (2020).

60. Zimmermann, J. et al. Gut-associated functions are favored during microbiome assembly across a major part of *C. elegans* life. mBio 15, e00012–24 (2024).

61. Buttin, G., Cohen, G. N., Monod, J. & Rickenberg, H. V. [Galactoside-permease of *Escherichia coli*]. Ann. Inst. Pasteur 91, 829–857 (1956).

62. Casale, A. et al. Complete genome sequence of *Escherichia coli* ML35. Genome Announc. 6, e00034–18 (2018).

63. Brenner, S. The genetics of *Caenorhabditis elegans*. Genetics 77, 71–94 (1974).

64. Arda, H. E. et al. Functional modularity of nuclear hormone receptors in a *Caenorhabditis elegans* metabolic gene regulatory network. Mol. Syst. Biol. 6, 367 (2010).

65. MacNeil, L. T., Watson, E., Arda, H. E., Zhu, L. J. & Walhout, A. J. M. Diet-induced developmental acceleration independent of TOR and insulin in *C. elegans*. Cell 153, 240–252 (2013).

66. Stiernagle, T. Maintenance of *C. elegans*. WormBook Online Rev. IC Elegansi Biol. 1–11 (2006) doi:10.1895/wormbook.1.101.1.

67. Jurkevitch, E. Isolation and classification of *Bdellovibrio* and like organisms. in Current Protocols in Microbiology (John Wiley & Sons, Inc., 2012).

68. Remy, O. et al. An optimized workflow to measure bacterial predation in microplates. STAR Protoc. 3, 101104 (2022).

69. Papkou, A. et al. The genomic basis of Red Queen dynamics during rapid reciprocal host–pathogen coevolution. Proc. Natl. Acad. Sci. 116, 923–928 (2019).

70. Davidov, Y., Friedjung, A. & Jurkevitch, E. Structure analysis of a soil community of predatory bacteria using culture-dependent and culture-independent methods reveals a hitherto undetected diversity of *Bdellovibrio*-and-like organisms. Environ. Microbiol. 8, 1667–1673 (2006).

71. Van Essche, M. et al. Development and performance of a quantitative PCR for the enumeration of Bdellovibrionaceae. Environ. Microbiol. Rep. 1, 228–233 (2009).

72. Caporaso, J. G. et al. Global patterns of 16S rRNA diversity at a depth of millions of sequences per sample. Proc. Natl. Acad. Sci. 108, 4516–4522 (2011).

73. Muyzer, G., de Waal, E. C. & Uitterlinden, A. G. Profiling of complex microbial populations by denaturing gradient gel electrophoresis analysis of polymerase chain reaction-amplified genes coding for 16S rRNA. Appl. Environ. Microbiol. 59, 695–700 (1993).

74. Martin, M. Cutadapt removes adapter sequences from high-throughput sequencing reads. EMBnet.journal 17, 10–12 (2011).

75. Callahan, B. J. et al. DADA2: High resolution sample inference from Illumina amplicon data. Nat. Methods 13, 581–583 (2016).

76. R Core Team. R: A language and environment for statistical computing. R Foundation for Statistical Computing (2024).

77. Camacho, C. et al. BLAST+: architecture and applications. BMC Bioinformatics 10, 421 (2009).

78. Kembel, S. W., Wu, M., Eisen, J. A. & Green, J. L. Incorporating 16S gene copy number information improves estimates of microbial diversity and abundance. PLOS Comput. Biol. 8, e1002743 (2012).

79. McMurdie, P. J. & Holmes, S. phyloseq: An R package for reproducible interactive analysis and graphics of microbiome census data. PLOS ONE 8, e61217 (2013).

80. Oksanen, J. et al. The vegan package. Community Ecol. Package 10, 631–637 (2007).

81. Brooks, M. E. et al. glmmTMB balances speed and flexibility among packages for zero-inflated generalized linear mixed modeling. R J. 9, 378–400 (2017).

82. Hartig, F. DHARMa: residual diagnostics for hierarchical (multi-level/mixed) regression models. CRAN Contrib. Packag. https://cir.nii.ac.jp/crid/1360583646752175744 (2016).

83. Lenth, R. V. emmeans: Estimated marginal means, aka least-squares means. 1.11.0 10.32614/CRAN.package.emmeans (2024).

84. Leo Lahti [Aut, C. microbiome. Bioconductor 10.18129/B9.BIOC.MICROBIOME (2017).

85. Kassambara, A. rstatix: Pipe-friendly framework for basic statistical tests. (2025).

86. Zimmermann, J., Kaleta, C. & Waschina, S. gapseq: informed prediction of bacterial metabolic pathways and reconstruction of accurate metabolic models. Genome Biol. 22, 81 (2021).

87. Zimmermann, J. et al. The functional repertoire contained within the native microbiota of the model nematode *Caenorhabditis elegans*. ISME J. 14, 26–38 (2020).

88. Lin, H. & Peddada, S. D. Multigroup analysis of compositions of microbiomes with covariate adjustments and repeated measures. Nat. Methods 21, 83–91 (2024).

89. Torchiano, M. effsize: Efficient effect size computation. (2020).

90. Champely, S., et al. pwr: Basic functions for power analysis. (2020).

91. Wickham, H. Ggplot2. (Springer International Publishing, Cham, 2016). doi:10.1007/978-3-319-24277-4.

92. Therneau, T. M. coxme: Mixed effects cox models. 2.2-22 10.32614/CRAN.package.coxme (2009).

93. Watson, E., MacNeil, L. T., Arda, H. E., Zhu, L. J. & Walhout, A. J. M. Integration of metabolic and gene regulatory networks modulates the *C. elegans* dietary response. Cell 153, 253–266 (2013).

94. Bito, T. et al. Vitamin B12 deficiency results in severe oxidative stress, leading to memory retention impairment in *Caenorhabditis elegans*. Redox Biol. 11, 21–29 (2017).

95. Virk, B. et al. Excessive folate synthesis limits lifespan in the *C. elegans*: *E. coli* aging model. BMC Biol. 10, 67 (2012).

96. Cabreiro, F. et al. Metformin retards aging in *C. elegans* by altering microbial folate and methionine metabolism. Cell 153, 228–239 (2013).

97. Annibal, A. et al. Regulation of the one carbon folate cycle as a shared metabolic signature of longevity. Nat. Commun. 12, 3486 (2021).

98. van der Goot, A. T. & Nollen, E. A. A. Tryptophan metabolism: entering the field of aging and age-related pathologies. Trends Mol. Med. 19, 336–344 (2013).

99. Qi, B. & Han, M. Microbial siderophore enterobactin promotes mitochondrial Iron uptake and development of the host via interaction with ATP synthase. Cell 175, 571–582.e11 (2018).

100. Feng, M., Gao, B., Ruiz, D., Garcia, L. R. & Sun, Q. Bacterial vitamin B6 is required for post-embryonic development in *C. elegans*. *Commun*. Biol. 7, 367 (2024).

101. Qi, B., Kniazeva, M. & Han, M. A vitamin-B2-sensing mechanism that regulates gut protease activity to impact animal’s food behavior and growth. eLife 6, e26243 (2017).

102. Bito, T., Matsunaga, Y., Yabuta, Y., Kawano, T. & Watanabe, F. Vitamin B12 deficiency in *Caenorhabditis elegans* results in loss of fertility, extended life cycle, and reduced lifespan. FEBS Open Bio 3, 112–117 (2013).

103. Revtovich, A. V., Lee, R. & Kirienko, N. V. Interplay between mitochondria and diet mediates pathogen and stress resistance in *Caenorhabditis elegans*. PLOS Genet. 15, e1008011 (2019).

104. Laranjeira, A. C., Berger, S., Kohlbrenner, T., Greter, N. R. & Hajnal, A. Nutritional vitamin B12 regulates RAS/MAPK-mediated cell fate decisions through one-carbon metabolism. Nat. Commun. 15, 8178 (2024).

105. Watson, E. et al. Metabolic network rewiring of propionate flux compensates vitamin B12 deficiency in *C. elegans*. eLife 5, e17670 (2016).

106. Bulcha, J. T. et al. A persistence detector for metabolic network rewiring in an animal. Cell Rep. 26, 460–468.e4 (2019).

107. Cheng, Y.-W., Liu, J. & Finkel, T. Mitohormesis. Cell Metab. 35, 1872–1886 (2023).

108. Ristow, M. & Schmeisser, K. Mitohormesis: Promoting health and lifespan by increased levels of Reactive Oxygen Species (ROS). Dose-Response Publ. Int. Hormesis Soc. 12, 288–341 (2014).

109. Na, H., Ponomarova, O., Giese, G. E. & Walhout, A. J. M. *C. elegans* MRP-5 exports vitamin B12 from mother to offspring to support embryonic development. Cell Rep. 22, 3126–3133 (2018).

110. Sivakala, K. K. et al. In vivo predation and modification of the Mediterranean fruit fly *Ceratitis capitata* (Wiedemann) gut microbiome by the bacterial predator *Bdellovibrio bacteriovorus*. J. Appl. Microbiol. 131, 2971–2980 (2021).

111. Gould, A. L. et al. Microbiome interactions shape host fitness. Proc. Natl. Acad. Sci. 115, E11951–E11960 (2018).

112. Henriques, S. F. et al. Metabolic cross-feeding in imbalanced diets allows gut microbes to improve reproduction and alter host behaviour. Nat. Commun. 11, 4236 (2020).

113. Wienhausen, G. et al. Ligand cross-feeding resolves bacterial vitamin B12 auxotrophies. Nature 629, 886–892 (2024).

114. Partridge, L., Piper, M. D. W. & Mair, W. Dietary restriction in *Drosophila*. Mech. Ageing Dev. 126, 938–950 (2005).

115. Greer, E. L. & Brunet, A. Different dietary restriction regimens extend lifespan by both independent and overlapping genetic pathways in *C. elegans*. Aging Cell 8, 113–127 (2009).

