## Supplementary figures for "Predatory bacteria impact *C. elegans* life-history traits by modulating microbiota community dynamics and thereby vitamin B12 availability"

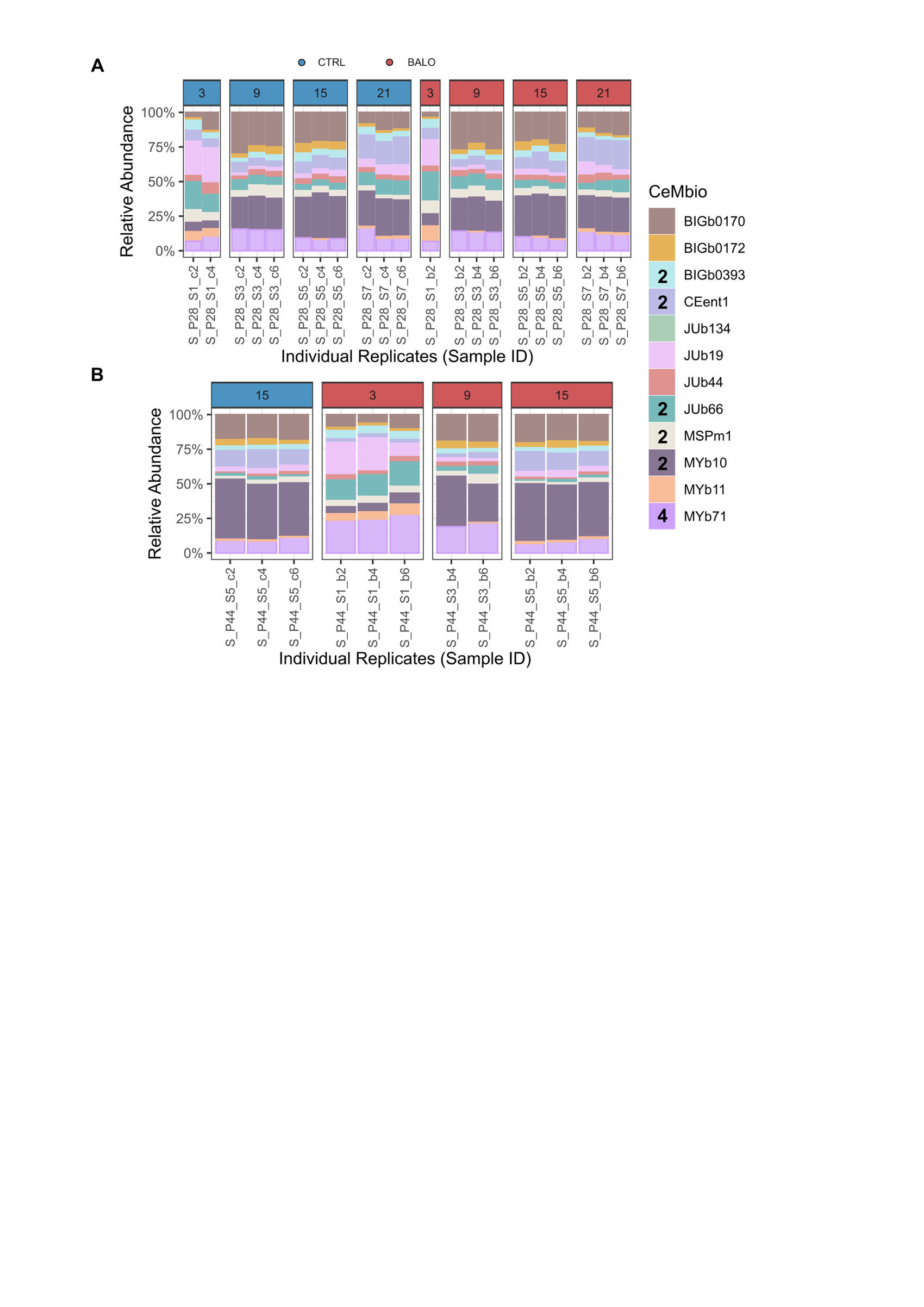

Fig. S1: Temporal dynamics of medium sample microbiome composition in response to BALO predation.
Relative abundances of community members within the host microbiome were determined via 16S amplicon sequencing and adjusted for 16S rRNA gene copy numbers. One bar represents the mean of up to 6 worm samples (technical replicates) for the (up to) 3 biological replicates that were used for the fitness assay. (A) Microbiome composition in the presence of the broad host-range BALO MYbb2. (B) Microbiome composition in the presence of the narrow host-range BALO MYbb4.

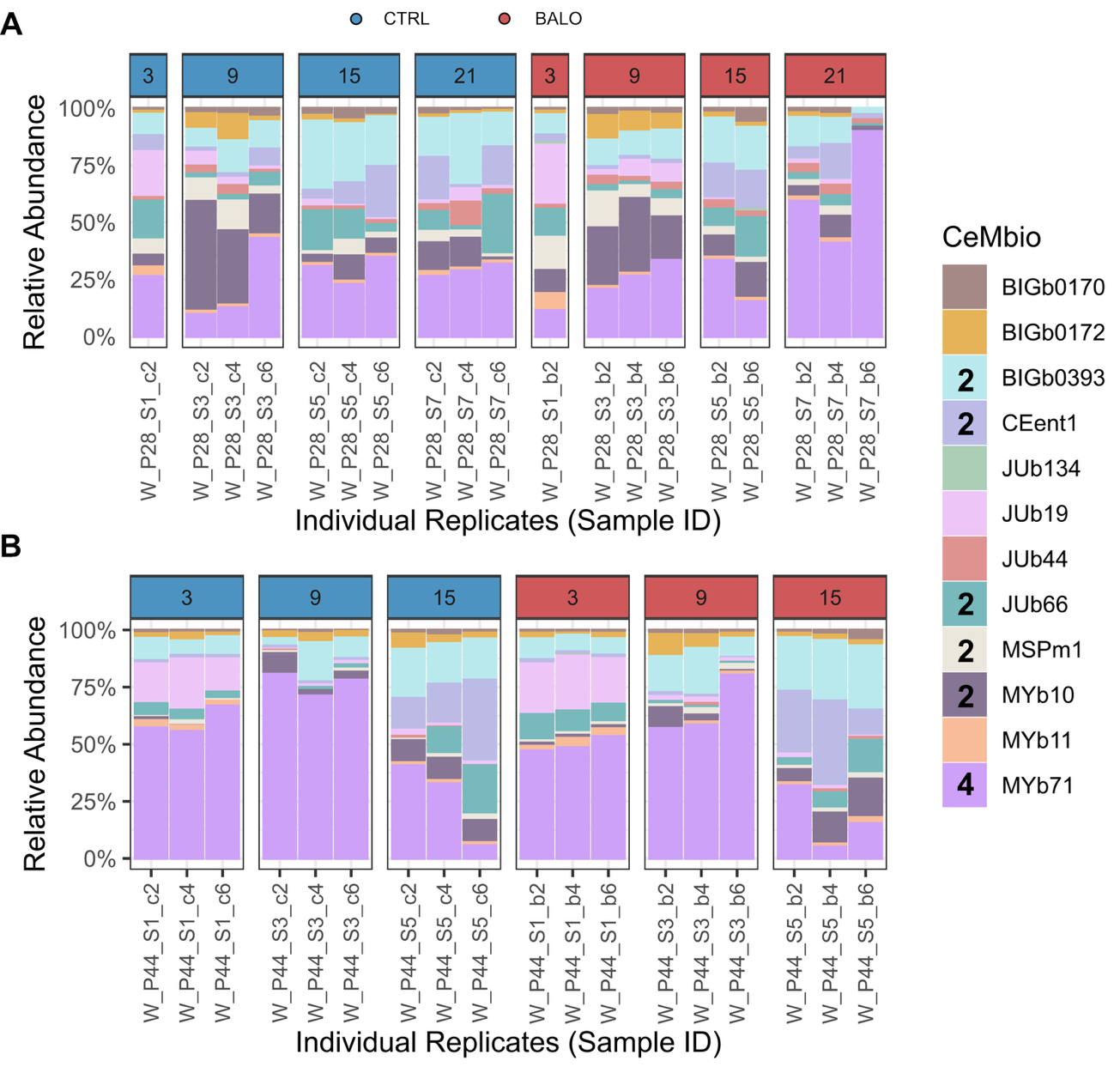

Fig. S2: Temporal dynamics of *C. elegans* microbiome composition in response to BALO predation.
Relative abundances of community members within the host microbiome were determined via 16S amplicon sequencing and adjusted for 16S rRNA gene copy numbers. One bar represents the mean of up to 6 worm samples (technical replicates) for the (up to) 3 biological replicates that were used for the fitness assay. (A) Microbiome composition in the presence of the broad host-range BALO MYbb2. (B) Microbiome composition in the presence of the narrow host-range BALO MYbb4.

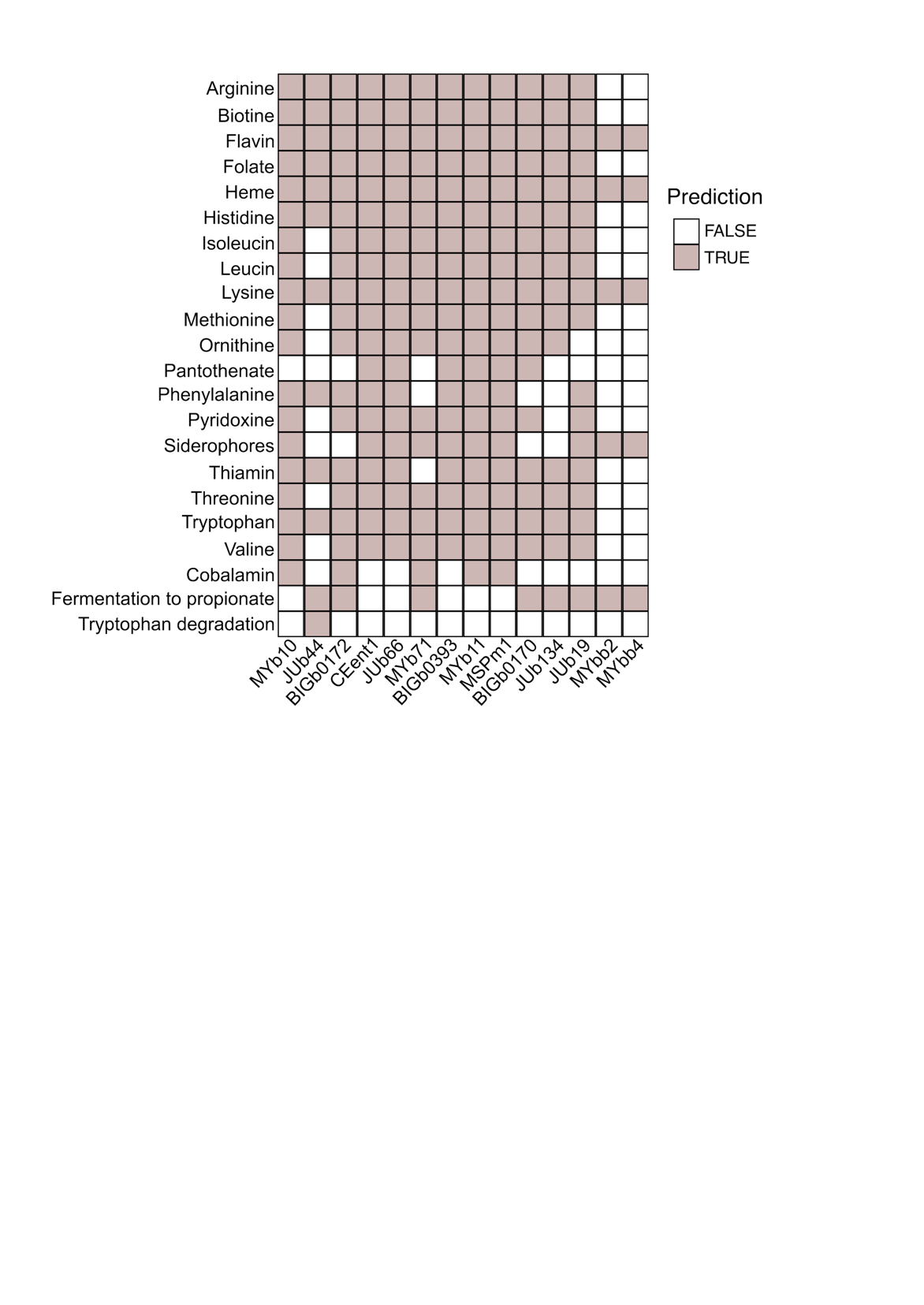

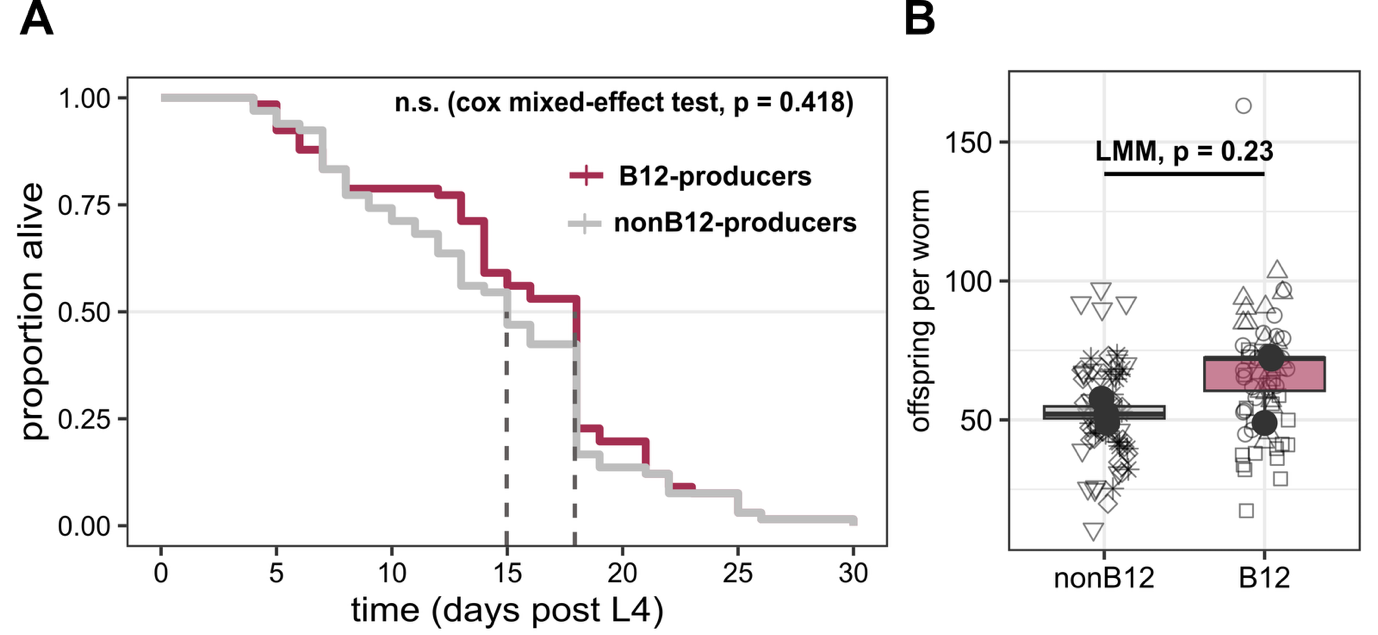

Fig. S3: Metabolic potential as inferred from whole genomes.

**Fig. S4: Host fitness outcomes of worms being exposed to bacteria producing vitamin B12 or non-vitamin B12-producing bacteria.** A: Kaplan-Meier curves show median lifespan in worms (Cox mixed-effects model). B: Mean offspring per worm across treatments, based on three biological replicates (n = 3). Unfilled symbols represent individual worms (technical replicates). Boxplots show replicate means. Groups were compared using a linear mixed effects model with biological replicate as random factor.

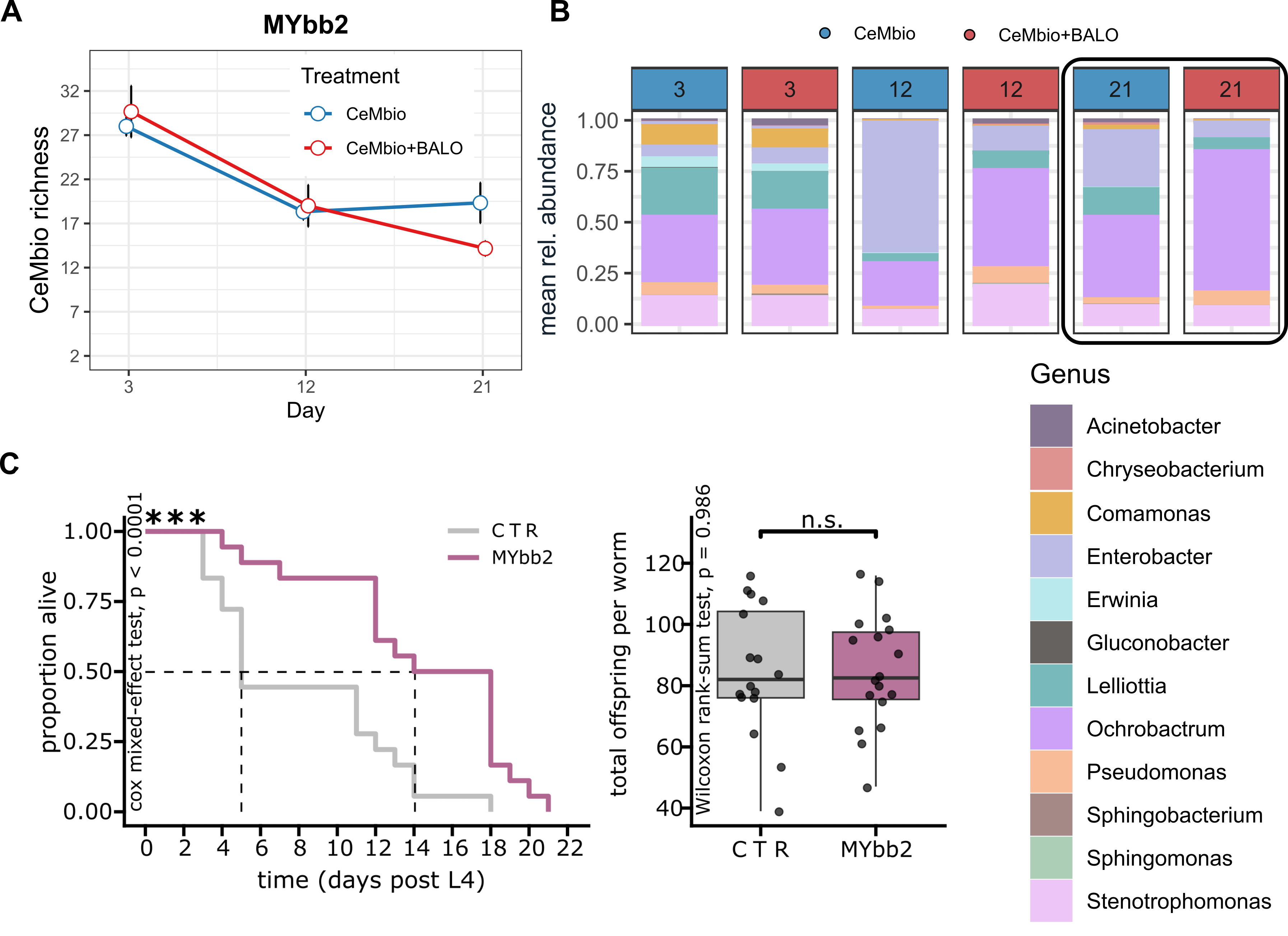

Fig. S5: Microbial community composition and host fitness following exposure to MYbb2 in the 43‑member CeMbio community. The experiment was conducted with 6 biological replicates (i.e., flasks) and 5 pooled worms per flask were analyzed. (A) Alpha‑diversity does not differ between worms treated with or without the broad host‑range BALO MYbb2. (B) Overall community structure is altered in the presence of MYbb2, with increased relative abundance of Ochrobactrum in treated worms. (C) Worms from the longitudinal diversity experiment exposed to MYbb2 exhibit a significantly extended lifespan, which persists despite a bleaching step performed after the diversity experiment. Kaplan-Meier curves (left) show median lifespan in MYbb2-treated worms (purple) compared to controls (grey; Cox mixed-effects model, p < 0.001). Mean offspring per worm across treatments (right), based on three biological replicates. Small dots represent individual worms (technical replicates; 6 per biological replicate).

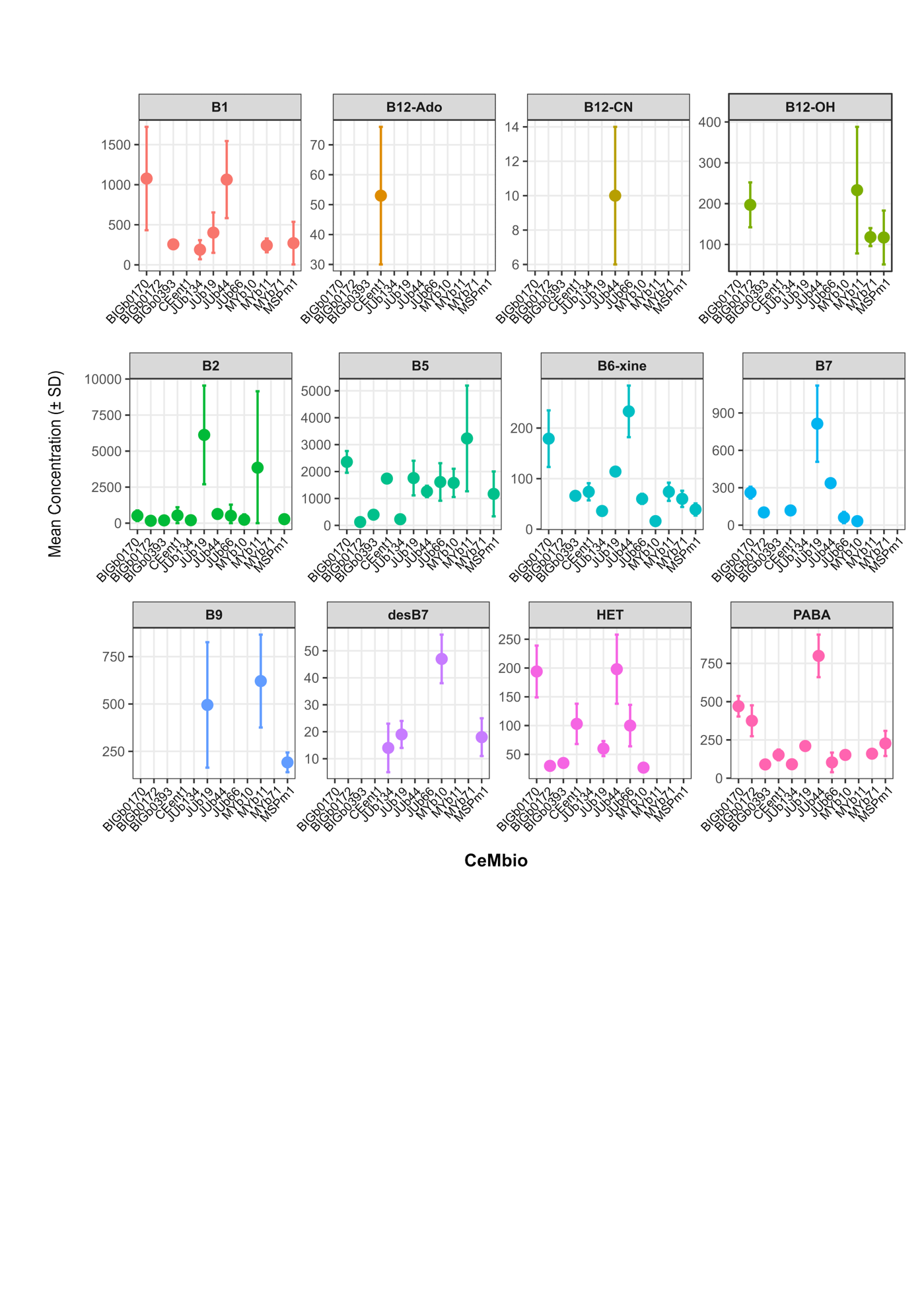

Fig. S6: Vitamin production by CeMbio strains. Amount (pmol of molecule extracted per gram of wet weight) of each vitamin per bacterial species. Only samples were at least 2 out of 3 biological replicates were above the limit of quantification are shown. Samples that fell below limit of quantification were removed.
